# RaDRI: A computational model for radiosensitisation by DNA double strand break repair inhibitors

**DOI:** 10.64898/2026.02.08.704735

**Authors:** Gib Bogle, Cho Rong Hong, Sophia F. O’Brien-Gortner, Barbara Lipert, Michael P. Hay, William R. Wilson

## Abstract

Repair of radiation-induced DNA double strand breaks (DSB) is a major contributor to radioresistance and an important target for tumour radiosensitisation. DNA-dependent protein kinase (DNA-PK) plays key roles in non-homologous end-joining (NHEJ), the dominant DSB repair pathway in human cells, and DNA-PK inhibitors (DNA-PKi) are highly effective radiosensitisers. However, many questions remain concerning tumour selectivity, mechanisms of enhancement of cell killing, interaction with other repair pathways and cell cycle checkpoints and the required duration of DNA-PK inhibition.

Here, we develop an agent-based computational model for Radiosensitisation by DSB Repair Inhibitors (RaDRI) with a level of complexity suitable for use in pharmacokinetic/pharmacodynamic models, and use it to investigate a potent and selective DNA-PKi, SN39536. RaDRI utilises analytical solutions for the spatial distribution of radiation-induced DSB, and their repair by NHEJ, from the Medras model (McMahon et al. Sci Rep 6:33290, 2016). Features include: (1) cell cycle progression and checkpoints are explicit; (2) probability of assignment of DSB to homologous recombination repair decreases with time post-replication, reflecting chromatin maturation, and is radiation dose-dependent; (3) Misjoining (ligation of ends from different DSBs), leading to chromosome aberrations, increases with time due to active DSB clustering.

The model is parameterised using flow cytometry and clonogenic survival datasets for low-LET irradiation of HCT116 cells, with and without the DNA-PKi. Clonogenic survival is computed as a function of the number of remaining DSBs and misjoins at mitosis. RaDRI demonstrates known radiobiological features including a near linear-quadratic dose dependence for killing by radiation, almost exclusively due to DSB misjoining, but predicts a distinct mechanism of radiosensitisation by SN39536 in which failure to resolve DSBs before mitosis becomes a significant driver of radiosensitisation. The model predicts that exposure to the DNA-PKi is required for ∼9 hours to achieve 90% of maximal radiosensitisation of DSB repair-proficient human cells in log-phase growth.

## 1. Introduction

Ionising radiation (IR) kills cells, and generates mutations, primarily through formation of DNA double strand breaks (DSB). The physical, chemical and biological processes involved are understood in detail over a wide range of timescales from the initial energy deposition events through to molecular, cellular and tissue responses over minutes to years. This deep mechanistic understanding, coupled with the importance of radiation therapy in cancer management, has encouraged development of a wide range of mathematical models that describe aspects of IR-induced biological effects^1-5^, including their modification by repair inhibitors^6^. However, prior work on modifiers of radiosensitivity predominantly model radiosensitisation via phenomenological survival curve modifications (e.g., α/β shifts in the linear-quadratic model)^7^ and few capture a contemporary understanding of the DSB repair pathways that influence resistance to IR.

The central role of DNA-dependent protein kinase (DNA-PK) in non-homologous end joining (NHEJ), the major DSB repair pathway in mammalian cells ^8^, has generated much interest in DNA-PK inhibitors (DNA-PKi) as radiosensitisers ^9,10^. NHEJ is available throughout the cell cycle and generally has high sequence fidelity in repairing DSBs ^11,12^ except when ends from different DSBs are ligated, which we refer to as misjoining, resulting in chromosome aberrations. The DNA-PK holoenzyme, comprising Ku70, Ku80 and the DNA-PK catalytic subunit (DNA-PKcs) has been the focus of elegant kinetic ^12,13^, structural ^14-16^ and mutational ^17,18^ studies that have recently converged to provide a detailed picture of the involvement of DNA-PK in NHEJ. Briefly, DNA-PK bridges DSB ends in two labile and interconvertible long-range synaptic complexes (the Ku-mediated dimer and XLF-dependent dimer) in which initial end processing can occur, followed by DNA-PKcs kinase-dependent transition of the XLF-dependent dimer to a short-range synaptic complex which lacks DNA-PKcs and mediates further end processing and end joining ^11,19^. However, while semi-mechanistic modelling of the DNA-PK inhibitor AZD7648 has been undertaken in a non-radiotherapy context ^20^, we are not aware of analogous models for radiosensitisation. Such models can potentially assist with questions such as (i) the relative contributions of unrepaired and misjoined DSBs to enhancement of radiation-induced cell killing ^21^; ^21^ whether a basal form of NHEJ can occur when DNA-PK kinase is fully inhibited; (iii) the role of potential backup DSB repair pathways in the absence of DNA-PK catalytic activity; (iv) the role of checkpoint delays in modulating radiosensitisation by DNA-PK inhibitors; and (v) the duration of DNA-PK inhibition required for radiosensitisation.

Our objective, here, is to develop a mechanistic model with complexity suitable for investigation of the pharmacology and mechanism of action of DNA-PKi as radiosensitisers. A particular motivation is to provide a computationally tractable single cell model that can be incorporated into agent-based models of large populations of tumour cells in order to explore the pharmacokinetics and pharmacodynamics (PK/PD) of DNA-PKi in tumour microenvironments. We have previously developed such “spatially resolved” PK/PD models for cytotoxic drugs and their hypoxia-activated prodrugs utilising a steady state approximation in which cell killing is a function of the area under the concentration-time curve (AUC) of the cytotoxic species at the target site ^21-25^, but this assumption is not applicable for DNA-PKi given that sensitivity to the inhibitors decreases as repair progresses. Thus we seek an agent-based model for radiosensitisation by DNA-PKi in which time as well as concentration dependence is explicit.

The model reported here (RaDRI, for <u>ra</u>diosensitisation by <u>D</u>NA repair inhibitors) builds on the Medras model developed by McMahon et al., ^26,27^ which we utilise for the formation and repair of IR-induced DSB and the relationship of unrepaired DSBs and chromosome aberrations to clonogenic cell killing. As explained in Section 3.1, we extend Medras through the following changes: (i) S-phase, which is typically the most radioresistant phase in the cell cycle, is included rather than representing the log-phase cell population as G1 and G2 phases only ^21^ ; ^21^ cell cycle progression is explicit, including effects of cell cycle checkpoints, the latter by further development of a computational model for ATM and ATR signalling in the G2 phase checkpoint ^28^ ; (iii) the probability of DSBs being assigned to HR decreases with time after DNA replication, reflecting chromatin maturation, and is suppressed at high radiation dose; (iv) the probability of misjoining via NHEJ increases with time post-irradiation, reflecting DSB mobility including active DSB clustering; (v) the role of DNA-PK as a determinant of NHEJ kinetics is explicit; (vi) although DNA damage at mitosis determines the probability of clonogenic cell killing, as in Medras, we model the surviving fraction of the population for clonogenic assays at a range of times post IR, not just at mitosis.

We estimate model parameters using clonogenic cell survival assays and changes in cell cycle distribution with HCT116 cells irradiated with and without SN39536 (Fig. 1), a potent and selective DNA-PKi which is an effective radiosensitiser in cell culture and tumour xenograft models ^29^. Although specification of the model requires a large number of parameters, reflecting the complexity of the biology, model fitting was constrained using literature information about ATM and ATR-dependence of cell cycle checkpoints, the kinetics of NHEJ and changes in the probability of HR and misrepair by NHEJ during progression through the cell cycle.

**Fig. 1.**
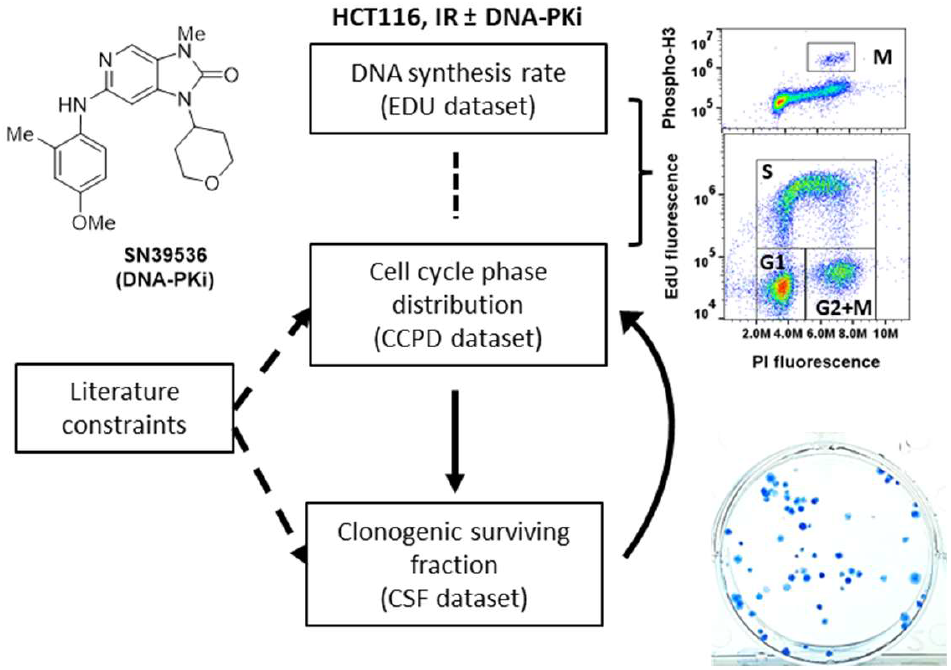
Workflow for RaDRI parameter estimation. Datasets describing effects of radiation with and without the DNA-PKi SN39536 were fitted using multiple cycles until convergence. Checkpoint parameters were estimated from the cell cycle phase distribution dataset (CCPD), determined using flow cytometry, and DNA damage, repair and lethality parameters from the clonogenic surviving fraction dataset (CSF).

## 2. Materials and methods

### 2.1 Cell lines and cell culture

Haploid HAP1 cells (catalogue C631) were purchased from Horizon Discovery and cultured in IMDM with 5% FBS (Morgate Biotech, Hamilton, New Zealand) until spontaneous reversion to diploidy. DLD1 cells with homozygous knockout of BRCA2 (DLD1/BRCA2^-/-^; catalogue HD105-007) and the isogenic parental DLD1-BRCA2^+/+^ line were also licensed from Horizon Discovery and were grown in DMEM with 5% FBS. The HCT116/54C cell line arose from a UT-SCC-54C culture that was contaminated with HCT116 cells which outgrew the cultures ^30^. HCT116/54C (henceforth HCT116) has been shown to closely resemble the ATCC line (HCT116/ATCC) in relation to ploidy, growth rate, colony morphology, plating efficiency, tumorigenicity, STR profile and mutational profile by exome sequencing ^30,31^ and, in the present study, radiosensitivity and radiosensitisation by SN39536 (Section 3.3). HCT116 and HCT116/ATCC were grown in MEM (ThermoFisher, 11095-MEM) supplemented 10% FBS, D-glucose to 4.5 g/L and 20 mM HEPES. All cell lines were maintained in log-phase growth without antibiotics for < 3 months from frozen stocks confirmed to be mycoplasma-negative by PlasmoTest (InvivoGen, San Diego, CA).

### 2.2 Repair inhibitors

SN39536 [6-((4-Methoxy-2-methylphenyl)amino)-3-methyl-1-(tetrahydro-2H-pyran-4-yl)-1,3-dihydro-2H-imidazo[4,5-c]pyridin-2-one; see Fig, 1] was synthesised as described ^29^. PolTheta inhibitor ART558 ^32^ was purchased from MedChemExpress (Monmouth Junction, NJ). Both compounds had purity > 99% by HPLC with diode array absorbance detection at 330 ± 50 nm and were stored as DMSO stock solutions at -20 °C.

### 2.3 Growth inhibition (IC_50_) assays

Repair inhibitors were added to 96 well plates seeded 2 h previously with 150 DLD1/BRCA2^+/+^ cells or 1250 DLD1/BRCA^-/-^ cells in 0.15 mL, using six 3-fold dilutions of each compound. Cultures were incubated for 7 days before determining cell number by staining with sulphorhodamine B and measurement of absorbance at 490 nm. IC_50_ values were estimated by 4 parameter logistic regression.

### 2.4 Irradiation

Cultures were initiated 5×10^4^ cells/mL in 24-well or P-60 dishes 24 h prior to irradiation, unless otherwise indicated in the figure legends. Dishes were irradiated at room temperature using a cobalt-60 teletherapy unit (Eldorado 78, Atomic Energy of Canada Ltd) at dose rates of 0.82-0.93 Gy/min, as determined by Fricke dosimetry, then returned to the 5% CO_2_ incubator at 37 °C for various growth times.

### 2.5 Cell cycle analysis

At various times after IR, S-phase cells were labelled with 100 µM 5-ethynyl-2’-deoxyuridine (EdU, Abcam) for 30 min. Cultures were then trypsinised and fixed with 70% ethanol at 4 °C overnight. Cells were permeabilised with PBST (PBS with 0.3% Tween 20) for 30 min at 4 °C and incorporated EdU was derivatised with 2 mM Alexa Fluor 647 (Thermo Fisher Scientific) using a click reaction (20 mM copper acetate and M Na-L-ascorbate in 20 mM Tris). Mitotic cells were detected by incubating in 0.5% normal goat serum (Abcam) in PBST for 1 h, staining with rabbit polyclonal a<u>nti-phospho-Histone H3 (Ser10) Antibody</u> (Merck, #06-570) for 1 h, washing in PBS and staining with Alexa Fluor 488-conjugated goat anti-rabbit IgG (Invitrogen, A32731) for 30 min. Cells were washed again in PBST and incubated in 0.25 mL PBS containing 20 µg/mL propidium iodide (Sigma) and 100 µg/mL RNAse (Serva) for 10 min before collecting data from 20,000 gated single cells with an Accuri C6 flow cytometer (BD Biosciences). Data were analysed using FlowJo software (v10).

### 2.6 Clonogenic cell survival assays

Cells from log-phase cultures were plated in 24-well or P-60 plates at 2.5×10^4^ – 10^5^ cells/mL and were irradiated 2-24 h later. SN39536 was added 1 h prior to IR and removed at later times by washing gently with pre-warmed medium. Clonogenic assays were performed 2-24 h after IR. For clonogenic assays, cultures were trypsinised, suspended in fresh medium, then cell densities were determined with an electronic particle counter (Z2 Coulter Counter, Beckman Coulter) and plated in triplicate at 50 – 5×10^4^ cells/well in 6 well plates. After growth for 10 days colonies were stained with methylene blue (2g/L in EtOH:H_2_O, 1:1 v/v) and colonies with >50 cells were counted manually to determine plating efficiencies (PE). Surviving fractions (SF) for radiation dose-response curves were calculated as the ratio PE(drug+radiation)/PE(drug only). Dose-response curves were fitted using the linear-quadratic model in GraphPad Prism (v10), or RaDRI. Sensitiser enhancement ratios (SER) were calculated as the ratio of radiation doses for isoeffect (e.g. SF = 10%) for no inhibitor/inhibitor.

### 2.7 Analysis of SN39536 in cell culture

Medium from HCT116/54C cultures in radiosensitisation experiments was stored at -20 °C then assayed by LC-MS/MS to determine SN39536 concentrations as previously ^29^.

### 2.8 Use of artificial intelligence

M365 Copilot (GPT-5 model) assisted with locating relevant literature, but had no role in writing the computer code or the text, or in parameter estimation or interpretation of the results.

### 2.9 Computational modelling

The RaDRI model is “agent-based”, meaning that a large number of cells (the agents) are simulated; each has a unique passage through the cell cycle to cell division. Initial cell cycle distribution is consistent with a log-phase steady state, and randomness influences a cell’s cycle progression (e.g. in the assignment of cycle duration). Statistics such as mean survival fraction are computed from the ensemble of simulated results, typically for 10,000 cells. The model is coded in Fortran 90. On a PC it runs under Windows, but for parameter estimation, which requires hundreds of model runs, the computer system of the New Zealand eScience Infrastructure Programme (NeSI) was used.

Parameter estimation was carried out employing two related programs, PEST_HP and CMAES_HP ^33^. These programs employ different methods to search for the best combination of a specified set of parameters to optimise the fit to experimental results. While PEST_HP employs “gradient” methods (based on numerical calculation of derivatives), CMAES_HP is a so-called “global” method, using stochastic generation of parameter values to explore the whole parameter space. In general, both methods were used sequentially, with CMAES_HP employed to check that PEST_HP had not become trapped in a local minimum.

The fit criterion was minimisation of the sum of squares of the weighted errors (objective function ϕ). The weight *w*(i) given to each measurement value *m*(i), which were log-transformed in the case of clonogenic surviving fractions and % of control for cell cycle phase distributions, was calculated as

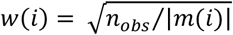

where *n*_*obs*_ is the number of observations (independent experiments).

The model parameters were estimated by fitting two experimental datasets describing effects of radiation on cell cycle phase distribution (CCPD dataset) and clonogenic surviving fraction (CSF dataset), each with and without the DNA-PKi. CCPD comprises the data shown in Fig. 8, and CSF in Figs 11 and 12. Parameter estimation proceeded iteratively, by cycling through the dataset fittings.

## 3. Results and Discussion

### 3.1 RaDRI model formulation: A computational model relating clonogenic cell killing to cell cycle progression and DSB repair

As in the Medras model, in RaDRI the probability of clonogenic survival is assessed at first mitosis after irradiation, based on the type and extent of genetic damage (numbers of unrepaired DSBs and misjoined chromosomes). Hence cell cycle checkpoints that facilitate completion of DSB repair prior to mitosis play a role and are incorporated explicitly. The model is parameterised by fitting experimental determinations of cell cycle perturbation and clonogenic cell survival of HCT116 cells following low LET (gamma) irradiation with or without the DNA-PK inhibitor SN39536. Given the complexity of the model, we also constrain many parameter values using literature data on the cell cycle dependence of DSB repair pathway choice and the kinetics and fidelity of repair on each pathway in human cells. Key features of the model are illustrated in Fig. 2 and explained more fully below.

**Table 2.** Cell cycle checkpoint parameters in RaDRI^a^.

| Parameter | Description | Value | Sensitivity of time to mitosis (h)<sup>b</sup> |  |  |  |  |  |
| --- | --- | --- | --- | --- | --- | --- | --- | --- |
|  |  |  | Mid-G1 |  | Mid-S |  | Mid-G2 |  |
|  |  |  | 6 Gy | 6 Gy +SN<sup>c</sup> | 6 Gy | 6 Gy +SN | 6 Gy | 6 Gy +SN |
| $k_{atm1G1}$ | Sensitivity of G1 slowdown to ATMact | 0.916 | -2.63, 8.33 | -1.06, 16.6 | - | - | - | - |
| $k_{atm2G1}$ | Sensitivity of G1 slowdown to ATMact | 0.429 | 0.53, -0.41 | 0.24, -0.18 | - | - | - | - |
| $k_{atm1G1d}$ | Sensitivity of G1 slowdown to ATMact in daughter cells | 030 | - | - | - | - | - | - |
| $k_{atm2G1d}$ | Sensitivity of G1 slowdown to ATMact in daughter cells | 0.0006 | - | - | - | - | - | - |
| $k_{atm1S}$ | Sensitivity of S phase slowdown to ATMact | 5.0 | -0.17, 0.16 | -0.22, 0.26 | -0.23, 0.27 | -0.08, 0.09 | - | - |
| $k_{atm2S}$ | Sensitivity of S phase slowdown to ATMact | 250 | 0.24, -0.12 | 0.38, -0.16 | 0.39, -0.17 | 0.14, 0.06 | - | - |
| $k_{mp1}$ | Rate constant for *ATMact* production from NHEJ-S | 1.32 | -0.51, 0.32 | -1.2, 0.82 | -0.19, 0.16 | 1.62, 1.06 | -1.15, 0.76 | -1.84, 1.16 |
| $k_{mp2}$ | Rate constant for *ATMact* production from HR | 0.685 | - | - | -0.04, 0.04 | -0.02, 0.02 | -0.11, 0.11 | -0.02, 0.01 |
| $k_{md}$ | Rate constant for *ATMact* decay | 8.81 | 0.52, -0.35 | 1.42, -0.94 | 0.38, -0.19 | 1.74, -1.27 | 1.38, -0.95 | 1.82, -1.38 |
| $K_{mmp}$ | $K_m$ for *ATMact* production | 14.9 | 0.32, -0.27 | 0.89, -0.67 | 0.19, -0.13 | 1.24, -0.99 | 0.91, -0.74 | 1.37, -1.14 |
| $K_{mmd}$ | $K_m$ for *ATMact* decay | 5.03 | -0.26, 0.2 | -0.58, 0.49 | -0.17, 0.15 | -0.68, 0.57 | -0.59, 0.49 | -0.71, 0.59 |
| $k_{rp,0}$ | Rate constant for *ATRact* production in the absence of DNA-PKi | 2.89 | - | - | -3.58, 2.79 | -0.23, 0.23 | -1.9, 2.99 | -0.14, 0.14 |
| $k_{rd}$ | Rate constant for *ATRact* decay | 2.91 | - | - | 5.04, -3.01 | 0.35, -0.17 | 4.93, -1.56 | 0.20, -0.10 |
| $K_{mrp}$ | $K_m$ for *ATRact* production | 497 | - | - | 3.85, -3.18 | -0.12, 0.12 | 4.50, -1.54 | 0.20, -0.11 |
| $K_{mrd}$ | $K_m$ for *ATR* decay | 0.112 | - | - | -0.2, 0.19 | -7.4, 3.6 | -0.82, 1.36 | -0.01, 0 |
| $k_{cc}$ | Rate constant for cell cycle (CC) activation | -<sup>d</sup> | - | - | - | - | - | - |
| $k_{ccmd}$ | Inhibition of CC activation by ATM | 75.7 | -0.05, 0.04 | -1.93, 1.51 | -0.20, 0.19 | -7.4, 3.6 | -5.82, 3.85 | -14.6, 6.24 |
| $k_{ccrd}$ | Inhibition of CC activation by ATR | 1.81 | - | - | -4.19, 3.75 | -0.23, 0.23 | -1.92, 3.28 | -0.14, 0.14 |
| $K_{mccp}$ | $K_m$ for CC activation | 0.0667 | 0, 0 | -0.02, 0.01 | 0.13, -0.13 | -0.05, 0.05 | 0.05, -0.05 | -0.14, 0.14 |
| $K_{mccmd}$ | $K_m$ for inhibition of CC activation by ATM | 647 | 0.07, -0.04 | 2.07, -1.44 | 0.28, -0.15 | 4.43, -5.26 | 2.93, -4.00 | 7.13, -10.3 |
| $K_{mccrd}$ | $K_m$ for inhibition of CC activation by ATR | 0.142 | - | - | 4.83, -2.84 | 0.18, -0.10 | 2.29, -0.77 | 0.01, -0.01 |
| $T_{lag}$ | Delay in onset of G1 slowdown (h) | 2 | - | - | - | - | - | - |
| $f_{thresh}$ | CCact threshold triggering entry to mitosis | 0.9 | - | - | - | - | - | - |

<sup>a</sup> All parameter values were estimated by fitting the CCPD dataset, except for $T_{lag}$ and $f_{thresh}$ which were
fixed. <sup>b</sup> Change in time to mitosis (changed value *minus* base value) when the parameter value is decreased by 1/3<sup>rd</sup> (first number) or increased by 1/3<sup>rd</sup> (second number). The base values for 6 Gy are 20.2 h for mid-G1, 23.1 h for mid-S and 15.3 h for mid-G2, and for 6 Gy + SN are 26.2 h, 25.0 h and 23.5 h, respectively. Light grey indicates moderate sensitivity (at least one value > 3 h) and dark grey indicates high sensitivity (at least one value > 10 h). <sup>c</sup> With 3 μM SN39536. <sup>d</sup> $k_{cc}$ takes a different value in each cell to equate its G2 phase duration, in the absence of DNA damage, to the initial phase duration assigned to that cell (see Section 3.1.5).

**Fig. 2.**
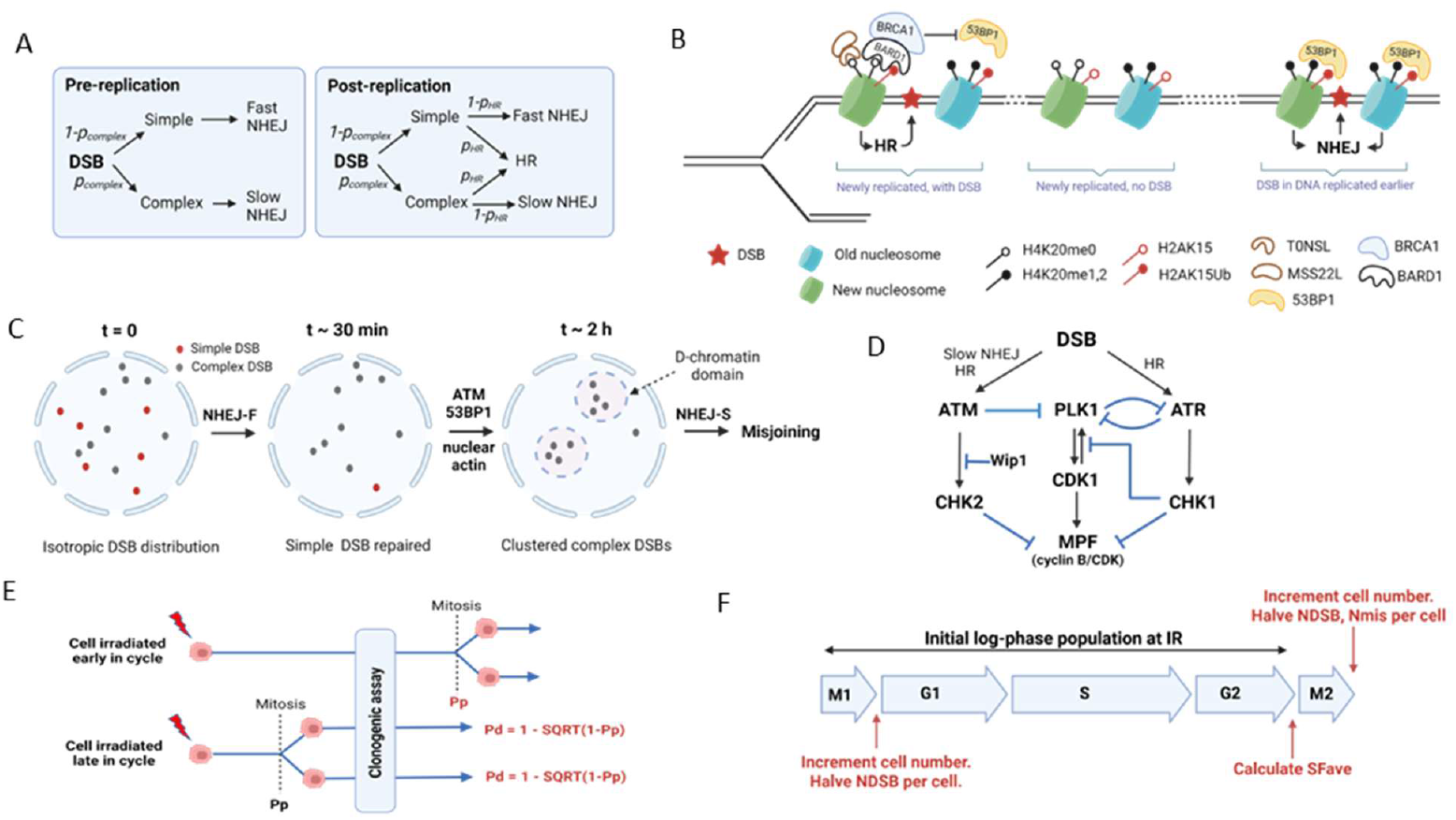
Features of the RaDRI computational model. A: Probabilities of assignment of DSBs to the three repair pathways represented in the model (fast NHEJ, slow NHEJ and HR) depend on ‘complexity’ of the DSB (see text) and whether the DSBs are generated in pre- or post-replication DNA. B: Histone epigenetic marks control access by the HR and NHEJ machinery in post-replication DNA; the H4K20me0 mark in newly recruited histones, required for HR, is progressively removed by methylation post-replication. C: DSBs induced by low LET radiation are initially isotropically distributed in the nucleus. Simple DSBs are rapidly repaired by fast NHEJ (NHEJ-F), while slow repair of complex DSBs by NHEJ-S is accompanied by active ATM- and 53BP1-mediated clustering of DSBs into damaged chromatin (D-chromatin) domains in G1-phase, enhancing misrepair by NHEJ and generating chromosome translocations, inversions and deletions (misjoins). D: Pathways controlling the G2-phase DNA damage checkpoint. Signalling cascades triggered by DSBs regulate activity of the mitosis promoting factor (MPF) through positive (black) and negative (blue) interactions. E: Clonogenic survival probability *p*_*survive*_ is calculated based on level of DNA damage at mitotic entry, but if the cell divides prior to clonogenic assay the survival probability of the daughter cells, *p*_*d*_, is calculated from the *p*_*survive*_ value for the parent cell (*p*_*p*_) at mitosis. F: Survival probability is determined at the next mitosis after irradiation, so that cells in mitosis at the time of IR (M1) are followed to the subsequent mitosis (M2) in order to capture the effects of DNA-PK inhibition in cells that survive IR in M1.

#### 3.1.1 Cell cycle phase distribution and progression

We assume for the untreated log-phase population that there is no cell loss and that the total cell cycle time of each individual cell, *T*_*C*_, is a log-normally distributed random variate with mean equal to the population doubling time *T*_*D*_. The mean phase durations, 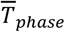, are calculated from the fraction of cells in each phase, *f*_*phase*_, as described previously for mitosis ^34^ and for the other phases ^35^. If the mean growth constant 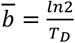 then:

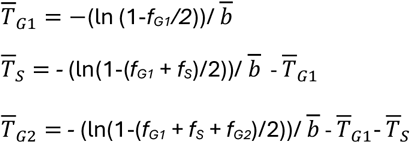

where *f*_*G2*_ is the measured fraction of cells with G2-phase DNA content after correction for the mitotic fraction.

Every cell, on creation (at model initialisation or after cell division), is assigned a cell cycle progression rate factor *f*_*pr*_ which modifies the duration of G1, S and G2 phases for that cell such that 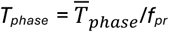). Individual values of *f*_*pr*_ are calculated to be consistent with *T*_*C*_ for each cell. Thus, if the duration of interphase for each cell is *T*_*inter*_ = *T*_*C*_ - *T*_*M*_, and the mean value for the population is 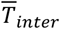, then 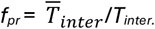. Variation in *T*_*M*_ is separately generated as a Gaussian random variate with mean value 0.56 h (section 3.2) and standard deviation 0.13 h, the latter from an Erlang distribution with k = 14, L = 28 which describes the distribution of mitotic durations in human cell lines ^36^.

After each cell has been assigned a cycle time and phase durations, cells are distributed across the cell cycle following Steel: the exponential probability density function of progress through the cell cycle, where *t* is time since the start of G1 phase, is:

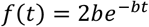

where *b* = *In*2/*T*_C_

This leads to the cumulative distribution function for the fraction of cells that have been in the cell cycle for less than time t:

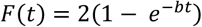

To generate *t* for a cell, a uniform (0,1) random variate R is generated, then:

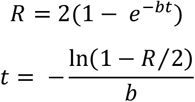

From *t* and the phase durations assigned to the cell, the cell’s initial phase and fractional progress through the phase are determined. In simulating cell cycle progression, a phase progress variable *P* is updated every time step (as explained in 3.1.5; in the case of checkpoint delay in G1 and S phases an additional factor fcp is included):

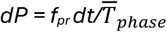

On entry to a phase, *P* is set to 0, and the cell makes the transition to the next phase when *P* = 1. In mitosis *f*_*pr*_ =1 and variation in M phase duration is determined as above. Progression in G2-phase is treated differently, with initiation of mitosis controlled by ATM and ATR signalling (Section 3.1.5).

#### 3.1.2 DSB formation, and assignment to repair pathways

We consider only two-ended DSBs generated by radiation, not single-ended DSBs that arise from stalled replication forks. Their initial number depends linearly on the radiation dose and the amount of DNA per cell, and hence on cell cycle phase. HCT116 is pseudodiploid ^31^; thus the number of DSBs created in G1 by a dose of *D* Gy, *N*_G1_ = *ψD* where *ψ* is a random number generated from a Poisson distribution with a mean of 35 DSB/Gy ^37^; if the fractional progress through S phase is *f*_*S*_, the total number of DSBs created is

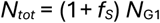

with *f*_*s*_ = 0 in G1 and 1 in G2.

Four different pathways repair radiation-induced DSBs in mammalian cells. The kinetics of the major pathway, NHEJ, depends in part on chemical features of the DSB (e.g. clustered base and sugar oxidations, hairpins) and on chromatin context, with a subset of “simple” DSBs undergoing much faster repair (NHEJ-F, half-time ca. 10 min) than “complex” lesions which are repaired by slow NHEJ (NHEJ-S, half-time ca. 2-3 h). Based on this kinetic distinction, pulsed field gel electrophoresis (PFGE) and γH2AX studies estimate the probability of DSBs being complex, *p*_*complex*_, as 0.43 ^26^ which we assume to be independent of dose and constant throughout the cell cycle. The NHEJ-S pathway has distinct molecular features, including activation of ATM and end-processing by the Artemis nuclease ^38^.

Homologous recombination (HR) repair requires extensive 5’ to 3’ resection by MRE11, EXO1 and the DNA2/BLM helicase, with early activation of ATM and subsequent recruitment of ATR at RPA/ATRIP-coated single stranded regions. HR effects relatively slow but high-fidelity repair of DSB in post-replication DNA utilising the DNA sequence in the sister chromatid as template during repair synthesis and is thus restricted to S/G2 phases in mammalian cells. PolTheta-mediated end-joining (TMEJ, aka alternate end-joining or microhomology-mediated end-joining (MMEJ)), and single-strand annealing ^39 39^, are slow highly error-prone DSB repair pathways. Both TMEJ and SSA, like HR, require 5’ to 3’ resection at DSBs which is initiated by S/G2-phase dependent CtIP activity exposing local microhomologies flanking the DSBs. Both are effectively subsumed within MMEJ in the Medras model, which allows this backup DSB repair throughout the cell cycle although it is now considered that TMEJ is initiated in S/G2 and completed during mitosis ^40-42^.

While the RaDRI formalism can readily accommodate TMEJ and SSA, in the present study we consider only three DSB repair pathways for reasons discussed below (Section 3.3): fast NHEJ (NHEJ-F) of simple DSBs, slow NHEJ (NHEJ-S) of complex DSBs and HR. We distinguish between DSBs generated in pre- and post-replication DNA, with numbers *N*_*pre*_ and *N*_*post*_, where:

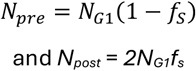

The probability of assignment to these three pathways is illustrated schematically for DSB generated in pre-replication and post-replication DNA in Fig. 2A. If *p*_*HR*_ is the probability of assignment to HR, and we make the simplifying assumption that simple and complex DSBs are equally accessible to HR, then the numbers of DSBs assigned to each pathway in pre- and post-replication DNA are given by:

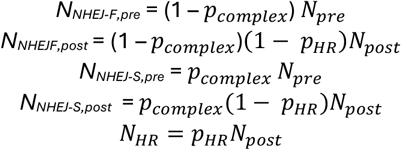

However, *p*_*HR*_ is a function not only of cell cycle stage but also of age (time since DNA replication) of sister chromatid sites at which DSBs are generated, and of radiation dose. The time dependence manifests as a decrease in pHR in late S/G2^43 44^ which is considered to be largely driven by chromatin maturation post-replication (Fig. 2B). Mechanisms include recruitment of unmethylated H4K20 to newly-replicated DNA which provides a histone mark that is read by ankyrin repeat domains (ARD) in the TONSL/MSS22L HR complex ^45^ and by the ARD in the BRCA1 binding partner BARD1 ^46^ to engage HR. This transient chromatin mark is extinguished by the cell cycle regulated monomethyltransferase SET8, which is expressed selectively on entry to G2 ^47,48^. The H4K20me0 post-replication mark operates in concert with a second chromatin mark generated by lysine 15 monoubiquinylation of H2A-type histones (H2AK15Ub) by RNF168 at DSB, which is read by a BUDR domain in BARD1 ^49^. These TONSL/MSS22L and multivalent BARD1 interactions enable assignment of DSBs to HR in newly replicated DNA, while BRCA1 recruited by BARD1 simultaneously blocks NHEJ by antagonising 53BP1 binding at DSBs (Fig. 2B). Thus we allow *p*_*HR*_ to decrease with time post-replication, resulting in lower values in G2 than in S-phase.

HR is a capacity-limited pathway ^50,51^ with *p*_*HR*_ suppressed at high radiation dose. In G2-phase, typical pHR values are 0.1-0.2 at low dose (1-2 Gy) ^52^ but PFGE studies at high dose indicate no repair defect in HR-defective cells ^53,54^. Similarly Mladenov et al. demonstrated that in G2 phase cells RAD51/γH2AX foci ratios fall with increasing radiation doses ^55^. In addition, gene conversion by HR in *I-Sce-*1 based reporter assays is strongly suppressed by IR ^55,56^. It is plausible that both the time-dependent and dose-dependent decreases in *p*_*HR*_ may have been evolutionarily selected to reduce risk of HR intermediates being present on entry to mitosis, potentially interfering with chromatid segregation.

To implement *p*_*HR*_ as a function of time and radiation dose in S- and G2-phases, we assume decay from initial level *p*_*HR*_ following a sigmoid function *f*_*decay*_(*t*):

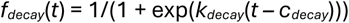

The decay is assumed to start after the cell has progressed through a specified fraction of S-phase, *f*_*S,decay*_ .

Dose suppression of *p*_*HR*_ is represented by a suppression factor *f*_*sup*_ where:

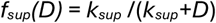

Thus overall:

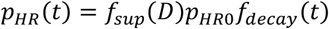

In other words, the time- and dose-variations of *p*_*HR*_ are characterized by the parameters *p*_*HRO*_, *k*_*sup*_, *f*_*S,decay*_, *k*_*decay*_ and *c*_*decay*_.

#### 3.1.3 Kinetics of DSB repair and misrepair

In each time step, of length *dt*, the number of unrepaired DSBs on the *j*th pathway is reduced by the factor 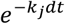, where *k*_*j*_ is the first order rate constant for repair on each pathway. For fast and slow NHEJ repair of simple and complex DSBs, based on McMahon’s analysis of PFGE data in repair-competent human cells ^27^, the respective values of *k*_*j*_ are *k*_*NHEJ-F*_ = 2.1 h^-1^ and *k*_*NHEJ-S*_ = 0.26 h^-1^. The kinetics of HR is less well defined. The slow phase of DSB repair in G2 after 2 Gy, monitored by γH2AX foci, has a rate constant ∼40% less than in G1_52_ but whether this reflects the contribution of HR is less clear. However, analysis of Rad51 foci after irradiation ^55,56^ indicates slower HR repair at high radiation dose, indicating that capacity limitions affect kinetics as well as *p*_*HR*_. We thus allow the rate constant for HR, *k*_*HR*,_ to decrease with dose from its maximum value at low dose, *k*_*HR,0*_ :

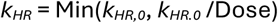

In relation to misrepair by NHEJ, we consider only the joining of incorrect DSB ends (misjoining) giving rise to chromosomal aberrations (translocations and intrachromosomal deletions/inversions); we ignore sequence changes at a DSB when the original ends are rejoined, which has little impact on clonogenic cell survival. Computation of misjoining utilises an analytical solution developed by McMahon for isotropically distributed DSBs from low LET radiation ^26^. Equations 12-15 in the latter paper provide the probability of misjoining, *P*_*mis*_, as a function of σ, a fitted scaling coefficient related to the rejoining range of NHEJ DSBs. However, because cell cycle progression is explicit in RaDRI we allow the nuclear radius (normalised to the G1 value) to increase by 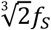 during S-phase.

The number of misjoined DSBs in a time step is then given by:

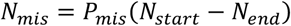

where *N*_*start*_ and *N*_*end*_ are the total numbers of NHEJ DSBs. The calculated number of pre-replication misjoins, given by *N*_*mis*_ (1-*f*_*S*_), is doubled at mitosis to reflect replication of misjoined chromosomes during S-phase.

We modify the treatment of misjoining by allowing movement of DBSs post-irradiation. In Medras the initial distribution of DSBs is isotropic and remains so with time, with the rejoining range specified by the parameter σ which is constant across the cell cycle and with time post-IR. However, there is much evidence for mobility of DSBs in the nucleus, which increases the probability of interaction between DSBs over time ^57,58^ and is consistent with evidence that the probability of chromosomal translocations is higher for repair by NHEJ-S than NHEJ-F ^59^. Further, there is clear evidence of ATM-dependent clustering of DSBs ^60^, particularly of slowly repaired DSBs in transcriptionally active genes during G1 ^61^. This clustering generates a distinct chromatin domain (D-chromatin) comprising activated DNA repair genes and in which chromosome translocation from nuclease-induced DSBs is enhanced ^62^. The damage-induced chromatin reorganisation, illustrated in Fig. 2C, is linked to a 53BP1-driven active phase separation process ^63-65^.

We accommodate this DSB mobility by allowing σ to increase with time post-IR. Thus:

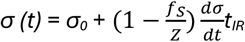

where *σ*^*0*^ is the initial value of *σ*. The rate of increase with time, *dσ/dt*, is specified by the input parameter *σ’* while *Z* controls the change in *σ’* across the cell cycle (suppression of clustering of post-replication DSBs). We assume that *σ*^*0*^ is constant through the cell cycle; although the cohesin complex contributes to radioresistance and DSB repair in G2 ^66^ and suppresses translocations in S-phase ^67^ there appears to be no difference in DSB mobility ^58^ or chromatin mobility ^68^ across the cell cycle with the exception of a transient mobilisation for ∼2 h after entry to G1 ^69^.

#### 3.1.4. Inhibition of NHEJ by DNA-PKi SN39536

In the model SN39536 inhibits the catalytic activity of DNA-PK (rate *DNAPKact*) which we assume has equal effect on both NHEJ-F and NHEJ-S. Inhibition of NHEJ is thus represented by reduction in the unihibited rate constants on both pathways (*k*_*NHEJ,F*_ and *k*_*NHEJ,S*_) as a function of *DNAPK*_*act*_ :

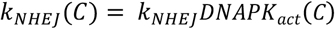

We represent the dependence of *DNAPK*_*act*_ on the extracellular concentration of SN39536, *C*, by a logistic function with upper and lower asymptotes at 1 and *DNAPK*_*act,min*_, respectively. Thus, as a fraction of the uninhibited activity:

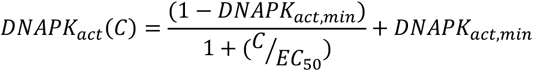

The concentration for 50% inhibition, *EC*_*50*_, and the lower asymptote *DNAPK*_*act,min*_ are estimated model parameters. We allow a non-zero value of *DNAPK*_*act,min*_to accommodate the possibility of non-canonical, DNA-PK-independent NHEJ as suggested by several lines of evidence ^11,70^ including that cells lacking DNA-PKcs are less radiosensitive than cells with deletion of core NHEJ factors ^71^.

#### 3.1.5. Radiation-induced cell cycle checkpoints

DSBs induce ‘emergency brakes’ that delay progression in each phase of the cell cycle, exploiting molecular pathways that overlap with the ‘intrinsic brakes’ that control cell cycle progression in undamaged cells ^72^. These delays are commonly referred to as checkpoints; we retain that widely used terminology although it carries the erroneous implication of arrest at specific points in the cell cycle. Both intrinsic and emergency brakes restrain cell cycle progression by controlling activity of cyclin/CDK complexes, but signalling from DSBs to the cell cycle machinery is orchestrated specifically by the ATM and ATR kinases. ATM/CHK2 signalling is an emergency brake only, while ATR/CHK1 signalling during DNA replication also acts as an intrinsic break to ensure correct cell cycle phase sequencing by suppressing mitotic kinase activity during S-phase ^73,74^.

Checkpoint delays in G2 are considered to have the greatest effects on radiation sensitivity because of their potential to increase time for DSB repair for cells irradiated close to mitosis, thus reducing mitotic catastrophe. Originally two separate G2-phase checkpoints were distinguished; an early but transient ATM-dependent checkpoint in cells irradiated late in G2, which is saturated at low radiation dose (∼ 1 Gy) and a late (slower) dose- and ATR-dependent checkpoint affecting cells irradiated in S- and early G2 phases ^75^. The transient nature of the ATM-dependent checkpoint was a longstanding puzzle because the activating S1981-ATM autophosphorylation triggered at DSBs occurs within minutes and is highly persistent ^76^, as is the case for other ATM autophosphorylations ^77^. Indeed even a 0.5 Gy dose (∼18 initial DSB) results in the S1981 phosphorylation of ∼ 50% of all ATM molecules in the cell ^76^, which was subsequently shown to result from a rapid ATM activation amplification loop via γH2AX, MDC1 and MRN that extends ATM recruitment ∼ 1 MB beyond the DSB ^39^.

However, it was subsequently shown that signalling to checkpoint pathways by ATM in these extended chromatin domains is more transient than ATM kinase activity at unrepaired DSBs or in the nucleoplasm ^28^. Specifically, the latter study used FRET reporters to demonstrate that in these chromatin domains ATM activity is rapidly terminated by chromatin-bound phosphatases such as Wip-1, enabling eventual PLK1 reactivation and mitotic entry. This finding is consistent with studies demonstrating that the G2 checkpoint is overcome before all DSBs are repaired ^78,79^. Phase progression in G2 is driven by positive feedback loops between mitotic kinases such as CDK1, CDK2, PLK1, Aurora A and its cofactor Bora ^80,81^ and their regulation by specific phosphatase complexes ^82^. This self-amplifying network is strongly suppressed by the kinase activities of ATM (e.g. via dephosphorylation of pT210-PLK1 by phosphatase PP2A/B55α) and ATR (e.g. via degradation of Bora), such that mitotic entry is controlled by opposing pro- and anti-mitotic signalling and does not necessarily require completion of repair ^83^. The main features of the pathways controlling the G2 checkpoint are illustrated schematically in Fig. 2D.

The Jaiswal study used a system of ODEs to model this control of mitotic entry in human G2-phase cells in response to the DSB-inducing antibiotic neocarzinostatin ^28^, integrating ATM and ATR signalling through their effects on the mitotic kinase network. We have built on that model, associating ATM and ATR signalling with resected DSBs assigned to HR, and ATM signalling to unresected complex DSBs assigned to NHEJ-S; we also allow the Michaelis constants controlling dependence on levels of DNA damage and ATM/ATR activity to take different values in each term of the ODEs.

ATM and ATR signalling to the checkpoints is controlled by the levels of ATM activation (*ATM*_*act*_) and ATR activation (*ATR*_*act*_) which in turn are determined by the unrepaired DSB count on each repair pathway. Note that these parameters are not equivalent to overall ATM and ATR phosphorylation or their kinase activity at DSBs, reflecting instead the overall status of the downstream checkpoint pathways. In the *ATM*_*act*_ differential equation the first term represents ATM activation by DSBs while second term represents phosphatase-dependent deactivation of ATM checkpoint signalling:

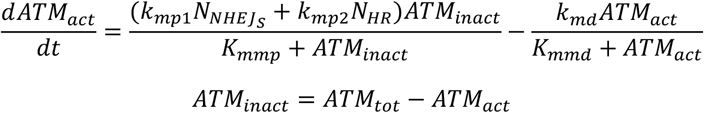

where *N*_*HR*_ and *N*_*NHEJs*_ are respectively the DSB counts assigned to HR, and complex DSB assigned to slow NHEJ repair, in each time step.

ATR activation occurs during HR in S and G2 phases. ATR checkpoint signalling, unlike ATM, is suppressed by high mitotic kinase activity represented by the cell cycle progression parameter *CC*_*act*_. Thus, the *ATR*_*act*_ differential equation is:

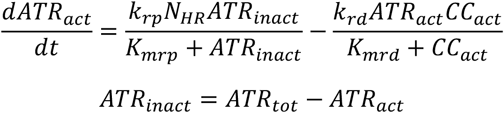

Although the above suffices in cells with basal DNA-PK catalytic activity, DNA-PK phosphorylates Ser4 and Ser8 in RPA32 to enhance ATRIP binding and thus ATR activation at resected DSBs ^84-86^. We represent this dependence of ATR signalling on DNA-PK activity (*DNAPK*_*act*_) as:

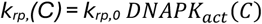

The differential equation for activation of cell cycle progression (*CC*_*act*_) includes a positive term for the mitotic kinase amplification loop, and negative terms for suppression of growth of *CC*_*act*_ signal via both ATM and ATR signalling:

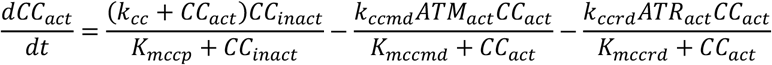

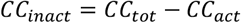

Entry to mitosis from G2-phase is triggered when *CC*_*act*_ rises to a threshold specified by *f*_*thres*_ *CC*_*tOt*_.

In the absence of DNA damage the G2 phase duration must equal the phase duration assigned to this cell (i.e. 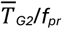), so it is necessary both to ensure that the parameters give rise to the correct phase duration in the no-damage situation and, for cells irradiated in G2, to initialise *CC*_*act*_ to the appropriate level. Since both *ATM*_*act*_ and *ATR*_*act*_ are zero when the cell is not irradiated, the duration of G2 is the time taken for *CC*_*act*_ to rise to the threshold level when only the first term in the above differential equation is active, i.e. the *CC*_*act*_ trajectory depends only on *k*_*cc*_ and *K*_*mccp*_. The approach adopted is to use the same value of *K*_*mccp*_ for all cells, and to determine *k*_*cc*_ for each cell such that the no-damage phase duration corresponds to the value assigned to that cell. In a similar way *CC*_*act*_ is initialized for each cell irradiated in G2. The no-damage differential equation for *CC*_*act*_ is integrated over the time the cell has spent in G2 to yield the initial value of *CC*_*act*_.

We utilise the same formalism for ATM activity in G1 and S-phases, and ATR activity in S-phase, with linkage of the apical kinase signals to checkpoint delay through a checkpoint factor *f*_*CP*_ that modifies the rate of progress through the phase:

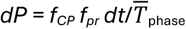

The ATM-dependent G1 checkpoint is controlled by slow, p53-mediated transcription generating the CDK2/cyclinE inhibitor p21, and separately by a rapid post-translational cascade that removes the CDC25A phosphatase; either pathway slows progression into S-phase. Given that the transcription-dependent pathway dominates in cells such as HCT116 with a functional p53 pathway, for simplicity we model *f*_*CP*_ as a simple empirical function of *ATM*_*act*_ with a time lag (*T*_*lag*_) to reflect the delay in the p21 transcriptional response, such that *f*_*cp*_ = 1 for time after IR <*T*_*lag*_, else:

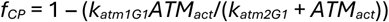

However, cell cycle delays in G1 are potentially different in post-mitotic cells, in part due to secondary DNA damage signalling triggered by micronuclei or breakage of dicentric chromosomes at cytokinesis. We therefore allow different G1 checkpoint parameters for cells that were irradiated in the previous cell cycle, with sensitivity of *f*_*CP*_ to *ATM*_*act*_ in these daughter cells defined by new parameters (*k*_*atm1G1d*_ and *k*_*atm2G1d*_) in the above equation.

The S-phase checkpoint is also primarily driven by ATM as originally demonstrated by radioresistant DNA synthesis in Ataxia Telangiectasia (ATM-null) cell lines ^87-89^. Hence in S-phase we model the checkpoint factor *f*_*CP*_ as for G1-phase but with parameters *k*_*atm1S*_ and *k*_*atm2s*_. ATR signalling plays a lesser role in the intra-S-phase checkpoint, so we ignore its effect on cell cycle progression in S-phase but we do allow ATR signaling during S-phase to set a non-zero value of *ATR*_*act*_ on entry to G2 and thus to enhance the G2 checkpoint for cells irradiated in S.

#### 3.1.6 Radiation-induced cell killing

In the present study we evaluate cell killing by clonogenic assay (CA) which effectively integrates all cell death pathways by evaluating long term proliferative survival of each cell. This approach potentially misses any cell loss (e.g. early apoptosis) occuring between the start of treatment and CA, but we show below (section 3.3) that there is no significant early cell loss up to 24 h after IR ± 3 μM SN39536 in our HCT116 cell line.

The probability of clonogenic cell survival for each cell, *P*_*survive*_, is computed on entry to first mitosis post-IR, except for cells irradiated in mitosis (M1) which are discussed below. The calculation is based on the number of misjoined chromosomes, *N*_*mis*_, and the number of unrepaired DSBs that were initially formed in pre- and post-replication DNA (*N*_*dsb1*_ and *N*_*dsb2*_,respectively) at mitosis. While misjoins are doubled during DNA replication in the model, rather than doubling *N*_*dsb1*_ post-replication we allow the two classes of DSB to have different effects on cell survival given that replication through DSB can generate more complex lesions. Thus:

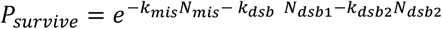

where model parameters *k*_*mis*_, *k*_*dsb*1_ and *k*_*dsb*2_ convey the influence on survival, at cell division, of chromosome aberrations, uprepaired DSB initially generated in pre-replication DNA and in post-replication DNA, respectively.

The simulation terminates when *P*_*survive*_ has been computed for all cells when they reach mitosis, whether mitosis occurs before or after CA. If no cells reached mitosis before CA the average survival probability *SF*_*ave*_ would be computed for the whole population as:

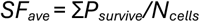

where *N*_*cells*_ is the total number of cells in the population at CA.

However, in the more general case *SF*_*ave*_ must be adjusted to account for daughter cells generated when parent cells reach mitosis before CA as illustrated in Fig. 2E. If the parent cell’s survival probability at mitosis is computed as *P*_*p*_, and the survival probability of each of its two daughters (both assumed equal) is *P*_*d*_, then the probability of no survival (i.e. reproductive death) of the parent is 1 – *P*_*p*_, which is equal to the probability of both daughters dying which is (1 – *P*_*d*_)^2^. Thus:

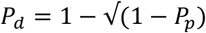

Therefore, in the calculation of Σ*P*_*survive*_ above, each cell reaching mitosis before CA is replaced by two cells with *P*_*surivive*_ = 1 – √(1-*P*_*p*_), and *N*_*cells*_ is incremented by 1.

Cells irradiated in mitosis (M1 in Fig. 2F) are highly radiosensitive because of chromatin compaction ^90^ and lack of DSB repair during mitosis other than by TMEJ ^40-42,91^. We represent the probability of clonogenic cell survival in M1 as:

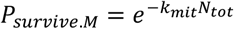

where *k*_*mit*_ has the value 0.023 which we estimate from the radiation survival curves for mitotic A2780 and OVCAR cells ^92^ assuming *N*_*tot*_ *=* 70 initial DSBs per Gy for these pseudodiploid cell lines. In the model, the daughters of cells that suffer lethal damage in M1 are followed until M2 (Fig. 2F) when probability of their survival is set to zero. The decision about lethality of damage in M1 is made by generating a uniform random variate *R* in the range (0,1); then the damage is lethal if *R* > *P*_*survive*.*M*_. For cells surviving irradiation in M1, *P*_*survive*_ is calculated from the number of DNA lesions of each kind at M2, as above. Following these cells to M2 is essential for modelling radiosensitisation by the DNA-PKi as their radiosensitivity is enhanced by NHEJ-driven chromosomal misjoining on entry to G1. Without accommodating their sensitisation due to misrepair in G1, cells irradiated in M1 become the most radioresistant at high DNA-PKi concentrations.

### 3.2 Cytokinetics of HCT116 cells

To establish cell cycle progression parameters for the model, we characterised the growth rate and cell cycle phase durations of our HCT116 cell line (Fig. 3). Log-phase cultures grew with a doubling time (*T*_*D*_) of 18.7 h (Fig. 3A) while flow cytometry using EdU pulse labelling, phospho-H3 and propidium iodide staining provided phase fractions (Fig. 3B) enabling calculation of mean phase durations (Fig. 3C). The estimated mean duration of mitosis, 0.56 h, was consistent with that for other human cell lines ^36^.

**Fig. 3.**
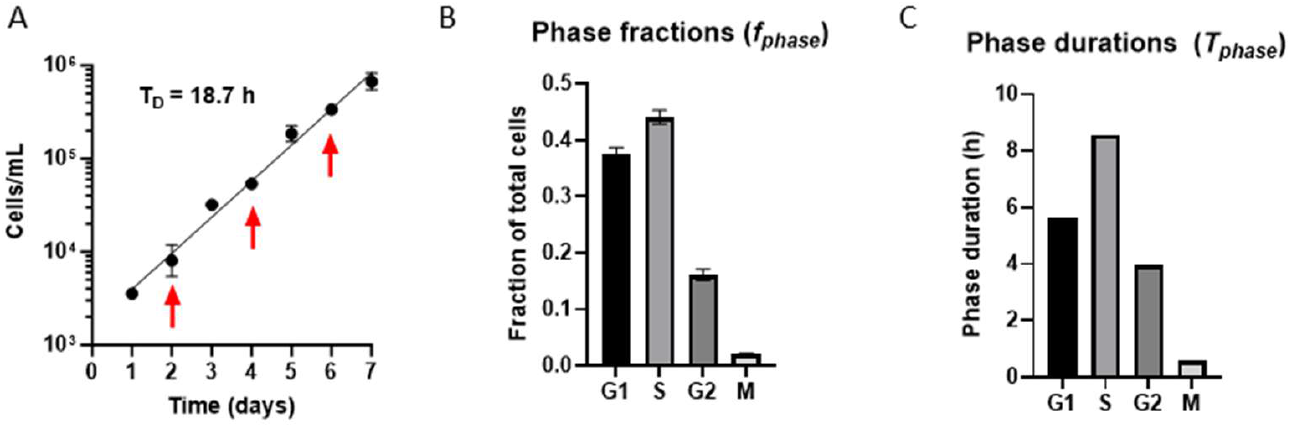
Growth kinetics, cell cycle phase distribution and mean phase durations of log-phase HCT116 cultures. A: Growth curve after seeding 10^4^ cells in 3 mL growth medium in 6-well plates (mean and SEM for 3 replicate cultures). Medium was replaced at the indicated times (arrows). B: Cell cycle phase fractions by flow cytometry (mean and SEM for 5 experiments). C: Mean phase durations calculated from phase fractions.

### 3.3 Radiosensitisation of HCT116 cells by DNA-PKi SN39536 and polTheta inhibitor ART558

We evaluated concentrations of SN39536 in cell culture as any compound instability would affect interpretation of time dependence of radiosensitisation. SN39536 at 3 µM was stable in HCT116 and HCT116/ATCC cultures for 25 h in a radiosensitisation experiment (Fig. 4) with final values 111 ± 9 and 112 ± 12 % (mean and SEM, N=3) of the initial concentration, respectively. The increases were not statistically significant and are consistent with a minor increase due to evaporation.

**Fig. 4.**
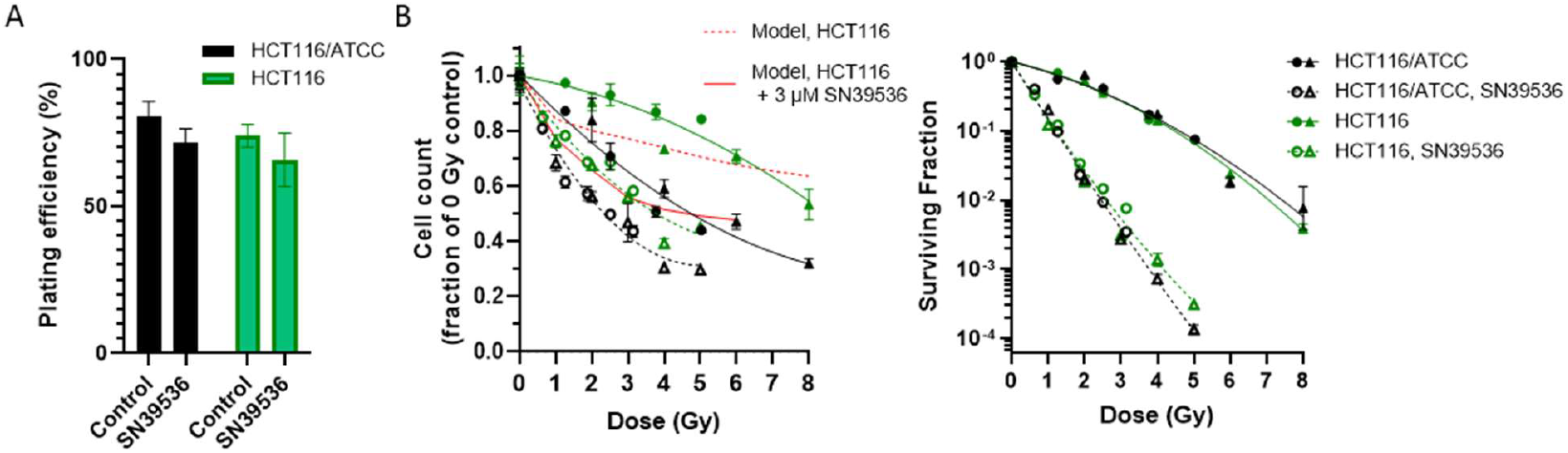
Radiosensitivity of HCT116 and HCT116/ATCC, and radiosensitisation by SN39536. Cells were plated at 5×10^4^/mL and irradiated one day later with or without SN39536 (3 µM) from 1 h before IR until clonogenic assay 24 h after IR. A: Plating efficiency for unirradiated controls. Values are mean and SEM for two experiments. B: Left, cell number at clonogenic assay, normalised to unirradiated values. Lines are quadratic fits. Red lines are RaDRI model predictions. Right, clonogenic surviving fraction. Values are mean and SEM for triplicate cultures in each experiment (shown with different symbols). Lines are linear-quadratic fits to the pooled values for the two experiments.

Radiosensitisation of HCT116 and HCT116/ATCC by 3 µM SN39536 was tested with cells exposed continuously to the drug for 1 h before and 24 h after IR at which time survival was tested by clonogenic assay (Fig. 4). This treat-then-plate experimental design facilitates control of drug exposure, but when plating is delayed (to allow inhibition of DNA-PK during the repair timescale) it becomes important to establish whether the drug alone causes cytotoxicity and whether there is inhibition of cell proliferation or cell loss prior to clonogenic assay. This 25 h exposure to the DNA-PKi at 3 μM had no significant effect on plating efficiencies (Fig. 4A; see also Fig. 11B). In both cell lines radiation reduced cell numbers at 24 h, with a greater effect in HCT116/ATCC; SN39536 increased this suppression in both lines (Fig. 4B, left panel). The model predictions of HCT116 cell number (red lines) are in reasonable agreement with measured values suggesting that cell cycle checkpoints (i.e. suppression of proliferation) rather than cell loss (e.g. by early apoptosis) are primarily responsible for the reduction of cell number at clonogenic assay: this is consistent with the lack of increase in the sub-G1 compartment by flow cytometry under these same conditions (data not shown). When measured by clonogenic assay 24 h after IR the two lines had indistinguishable radiosensitivity and were equally radiosensitised by SN39536 (Fig. 4B, right panel), with SER_10_ values of 3.81 ± 0.29 (HCT116) and 3.83 ± 0.06 (HCT116/ATCC).

Given that TMEJ potentially acts as a backup repair pathway when NHEJ is inhibited ^40-42,93,94^, we then evaluated radiosensitisation of HCT116 by SN39536 and the polTheta inhibitor ART558, individually and in combination (Fig. 5). We confirmed the expected synthetic lethal interaction between ART558 and BRCA2 deficiency using a DLD-1 isogenic pair (Fig. 5A), although the differential was less than reported previously in these cell lines ^32^ and much lower than the 50-fold differential for olaparib with these cell lines determined in our lab using the same methodology ^95^. In contrast, SN39536 did not demonstrate a significant interaction with *BRCA2* mutation (Fig. 5A).

**Fig. 5.**
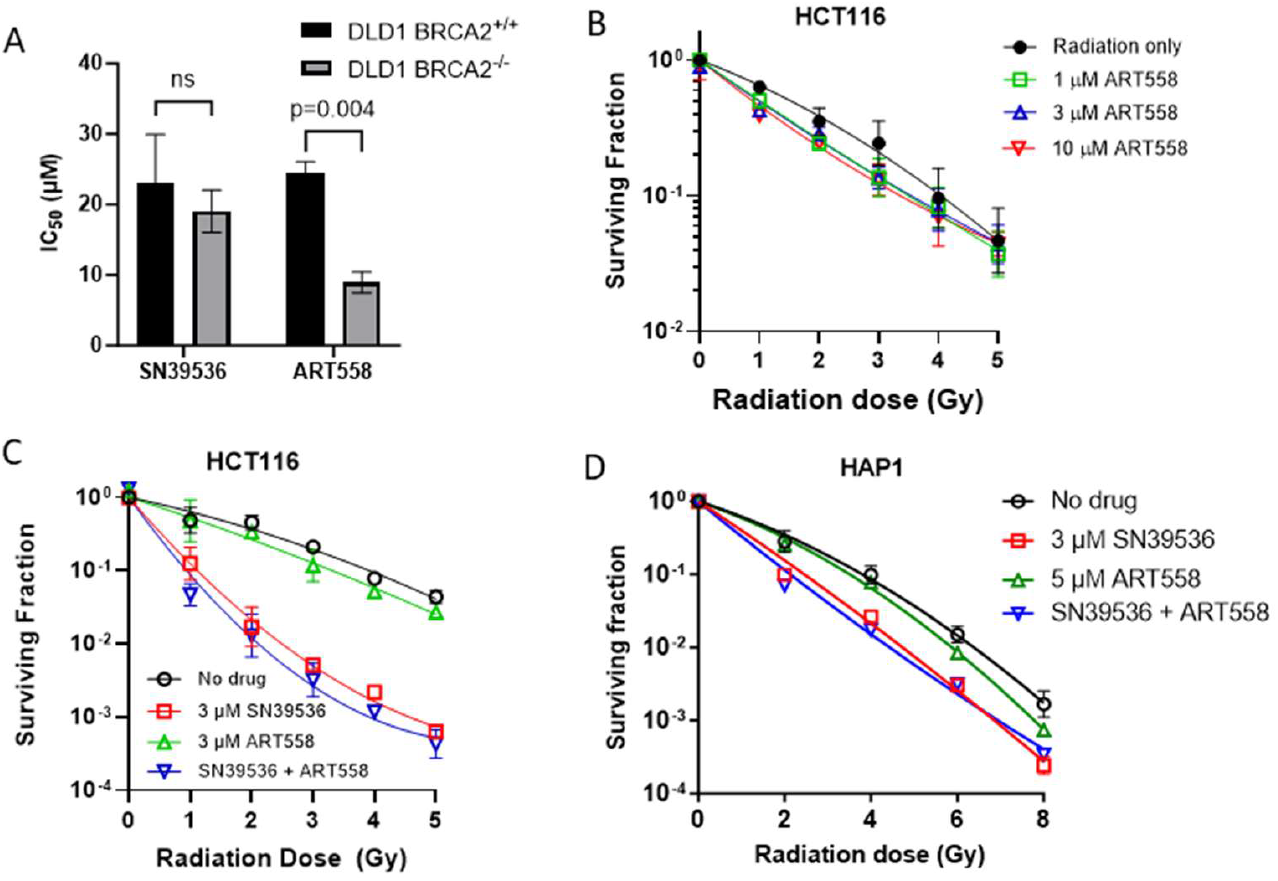
BRCA2-dependence of cytotoxicity of SN39536 and pol-theta inhibitor ART558, and radiosensitisation by each alone and in combination. A: Growth inhibition of log-phase monolayers of wild-type DLD1 cells and an isogenic *BCRA1* knockout line after continuous exposure to the compounds for 7 days. Values are mean and SEM for 3 experiments. B,C: Radiosensitisation of log-phase HCT116 cells to ART558 (B) or both compounds individually and together at 3 µM (C) for 1 h before IR and then for 18 h until clonogenic assay. D: Radiosensitisation of HAP1 cells by 5 µM ART558, 3 µM SN39536 and their combination with drug exposure 1 h before and for 18 h until clonogenic assay. Values are mean and SEM for 2 experiments in B-D, and lines are fits to the L-Q model.

ART558 provided modest radiosensitisation of HCT116 over the concentration range 1-10 µM (Fig. 5B) with an SER_10_ of 1.23 ±0.01 at 10 µM. At 3 µM, ART558 did not enhance radiosensitisation by SN39536 (Fig. 5C); SER_10_ values were 1.29 ± 0.01 for ART558 alone, 3.52 ± 0.19 for SN39536 alone and 4.20 ± 0.19 for the combination; the ratio of SER_10_ values for SN39536 with or without ART558 was 1.17 ± 0.02 (no greater than the SER_10_ for ART558 alone) suggesting only a minor additive effect. Similar results were obtained with the HAP1 cell line in which 5 µM ART558 again provided only marginal radiosensitisation and did not increase sensitisation by the SN39536 (Fig. 5D). Given the minor effect of ART558 on radiosensitivity, and lack of enhancement of sensitisation by the DNA-PKi, we have not included TMEJ repair in the current iteration of RaDRI.

### 3.4 RaDRI model

We parameterised the RaDRI model by iterative fitting to two datasets (cell cycle phase distributions in CCPD and clonogenic surviving fractions in CSF) with radiation alone and in combination with SN39536 as described in Section 2.9. The final values of the parameters that control initial DSB yields, repair pathway commitment, repair kinetics and lethality of the DNA lesions at mitosis are shown in Table 1. Some parameters were fixed based on literature values, some of the parameters to which the model had low sensitivity were assigned in order to represent known radiobiology, and some were estimated by unconstrained fitting. The assigned parameters were not routinely estimated through the CCPD/CSF fitting cycles but were in some cases estimated with tight bounds late in the process to optimise fits.

#### 3.4.1 Commitment of initial radiation-induced DSBs to repair pathways, and repair kinetics

Initial DSBs per cell vary across the cell cycle in proportion to DNA content and radiation dose, and are allocated to repair pathways according to complexity of the lesions and availability of HR as described in Section 3.1.2 and illustrated for the final parameter set in Fig. 7A. Given that sensitivity of cell killing to the parameters controlling pHR (*p*_*HRO*_, *k*_*sup*_, *f*_*S,decay*_, *k*_*decay*_ and *c*_*decay*_) was very low (see sensitivity of *D*_*10*_ in Table 1; full survival curves are shown in Fig. S1), the HR parameters were assigned to provide features consistent with literature rather than being estimated from the CSF data. The key features include a high probability of HR immediately after IR in post-replication DNA during S-phase (*p*_*HR,0*_ = 1), decay of pHR from late-S (*k*_*fS,decay*_ = 0.8) declining to an average of 0.125 in G2 after 2 Gy and to 0.049 after 6 Gy (Fig. 6A) as a result of suppression of HR at high dose. The DSB repair rates are dose-independent for NHEJ and dose-dependent for HR (Fig. 6B). The change in pHR with time post-replication, as a result of chromatin maturation, is illustrated in Fig. 6C.

**Fig. 6.**
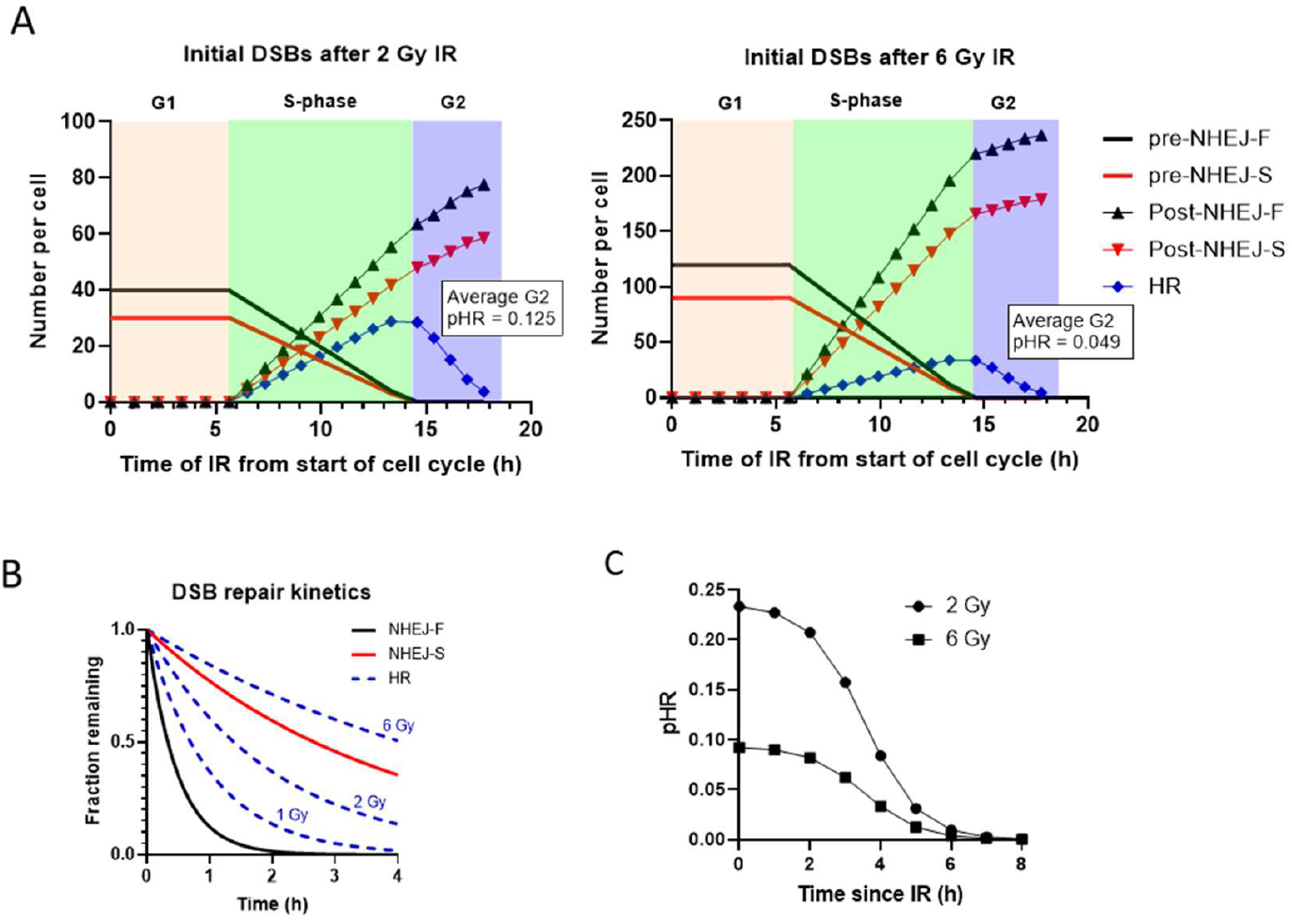
RaDRI model allocation of initial radiation-induced DSBs to repair pathways as a function of cell cycle stage, and DSB repair kinetics. A: DSBs in pre-replication DNA are allocated to fast NHEJ (pre-NHEJ-F) or slow NHEJ (pre-NHEJ-S) depending on lesion complexity. In post-replication DNA, fast NHEJ (post-NHEJ-F), slow NHEJ (post-NHEJ-S) and HR are all available, with probability of HR decreasing with time and radiation dose. B: Repair kinetics on the NHEJ pathways (independent of dose) and HR repair kinetics illustrated at 3 dose levels. C: Decay of pHR with time after irradiation during S-phase, for DSBs generated immediately after replication (p_HR,o_ = 1).

#### 3.4.2. Measurement and RaDRI modelling of radiation-induced checkpoint delays

The behaviour of the core checkpoint model, based on the Jaiswal formulation of ATM and ATR signalling from unrepaired DSBs ^28^, is illustrated for cells irradiated at the G2/S border, using the final checkpoint and DSB repair parameter values in Tables 1 and 2 (Fig. 7). ATM signalling (*ATM*_*act*_) is earlier and more transient than for ATR (*ATR*_*act*_), and is prolonged in combination with 3 μM SN39536 as a result of delayed NHEJ-S repair, while there is essentially no ATR signalling as a consequence of the dependence of ATR activation on DNA-PK activity in the model. For radiation alone, *ATR*_*act*_ is the major determinant of entry to mitosis (Fig. 7A,B) while the absence of ATR signalling when DNA-PK is strongly inhibited results in *ATM*_*act*_ controlling entry to mitosis (Fig. 7C,D).

**Table 1.** DNA damage, repair and radiosensitivity parameters estimated by RaDRI using the SFALL dataset^a^.

| Parameter | Description | Units | Source<sup>b</sup> | Parameter value | Sensitivity of D<sub>10</sub> (Gy)<sup>c</sup> |  |
| --- | --- | --- | --- | --- | --- | --- |
| | | | | | IR only | IR + 3 μM SN39536 |
| **DNA damage parameters** |  |  |  |  |  |  |
| $\psi$ | Initial number of DSBs in cells irradiated in G1 phase | Gy<sup>-1</sup> | Fixed | 35<sup>d</sup> | - | - |
| $p_{\text{complex}}$ | Probability that a DSB is complex | - | Fixed | 0.43 | - | - |
| **DNA repair parameters** |  |  |  |  |  |  |
| $k_{\text{NHEJ-F}}$ | Rate constant for fast NHEJ (NHEJ-F) | h<sup>-1</sup> | Fixed | 2.08 | - | - |
| $k_{\text{NHEJ-S}}$ | Rate constant for slow NHEJ (NHEJ-S) | h<sup>-1</sup> | Fixed | 0.26 | - | - |
| $k_{\text{HR},0}$ | Rate constant for HR at low radiation dose | h<sup>-1</sup> | Assig. | 1.0 | 0.08, -0.09 | 0.02, 0 |
| $p_{\text{HR},0}$ | Probability that a DSB formed in newly replicated DNA is assigned to HR | - | Assig. | 1.0 | -0.24, - | -0.04, - |
| $k_{\text{decay}}$ | Parameter controlling decay of $p_{\text{HR}}$ post-replication | - | Assig. | 1.29 | -0.03, 0 | -0.01, 0.01 |
| $C_{\text{decay}}$ | Parameter controlling decay of $p_{\text{HR}}$ post-replication | - | Assig. | 3.54 | -0.10, 0.07 | -0.03, 0.01 |
| $f_{\text{s,decay}}$ | Fraction of $f_{\text{s}}$ at which decay of $p_{\text{HR}}$ starts | - | Assig. | 0.8 | -0.18, 0.10 | -0.05, 0.03 |
| $k_{\text{sup}}$ | Factor controlling suppression of $p_{\text{HR}}$ with radiation dose | - | Assig. | 0.617 | -0.21, 0.16 | -0.04, 0.04 |
| $\sigma$ | Characteristic rejoining range of DSBs misrepaired by NHEJ | Fraction of $R^e$ | Est. | 0.0531 | 1.73, -1.05 | 0.20, -0.16 |
| $\sigma'$ | Rate of increase of $\sigma$ post-irradiation | h<sup>-1</sup> | Est. | 0.0326 | 0.71, -0.55 | 0.44, -0.15 |
| $Z$ | Controls suppression of misjoining in post-replication DNA | - | Est. | 2 | 0.48, -0.22 | 0.42, -0.09 |
| **DNA-PK inhibition parameters** |  |  |  |  |  |  |
| $EC_{50}$ | Concentration (in extracellular medium) for 50% inhibition of DNAPK<sub>act</sub> by SN39536 | μM | Est. | 0.232 | 0, 0 | -0.09, 0.08 |
| $N$ | Hill coefficient for inhibition of DNAPK<sub>act</sub> by SN39536 | - | Fixed | 1.0 | - | - |
| $DNAPK_{\text{act,min}}$ | Asymptote of DNAPK<sub>act</sub> at high SN39536 concentration | - | Est. | 0.09 | 0, 0 | -0.12, 0.11 |
| **Lethality parameters** |  |  |  |  |  |  |
| $k_{\text{mis}}$ | Coefficient for effect of misjoins on probability of survival | - | Est. | 0.391 | 1.00, -0.58 | 0.47, -0.20 |
| $k_{\text{dsb1}}$ | Coefficient for effect of DSBs in pre-replication DNA on probability of survival | - | Est. | 0.152 | 0.02, -0.02 | 0.19, -0.16 |
| $k_{dsb2}$ | Coefficient for effect of DSBs in post-replication DNA on probability of survival | - | Est. | 0.0677 | 0.05, -0.04 | 0.18, -0.14 |
| $k_{mit}$ | Coefficient for effect of DSBs generated in mitotic cells on probability of survival | - | Fixed | 0.023 | - | - |

<sup>a</sup> SFALL comprises clonogenic surviving fractions for radiation with and without SN39536 at a range of concentrations and drug exposure times. <sup>b</sup>Parameter values were fixed from literature, assigned (assign.) to represent known radiobiology or estimated (Est.). <sup>c</sup> Change in dose for 10% clonogenic survival ( $D_{10}$ changed value *minus* $D_{10}$ base value) when the parameter value is decreased by $1/3^{rd}$ (first number) or increased by $1/3^{rd}$ (second number), where the base value of $D_{10}$ is 4.12 Gy for IR alone and 1.38 Gy for IR + 3 μM SN39536. Parameters to which the model is most sensitive (>0.3 log change) are highlighted in grey. <sup>d</sup>Mean value (Poisson distribution). <sup>e</sup>Normalised G1 phase nuclear radius.

**Fig. 7.**
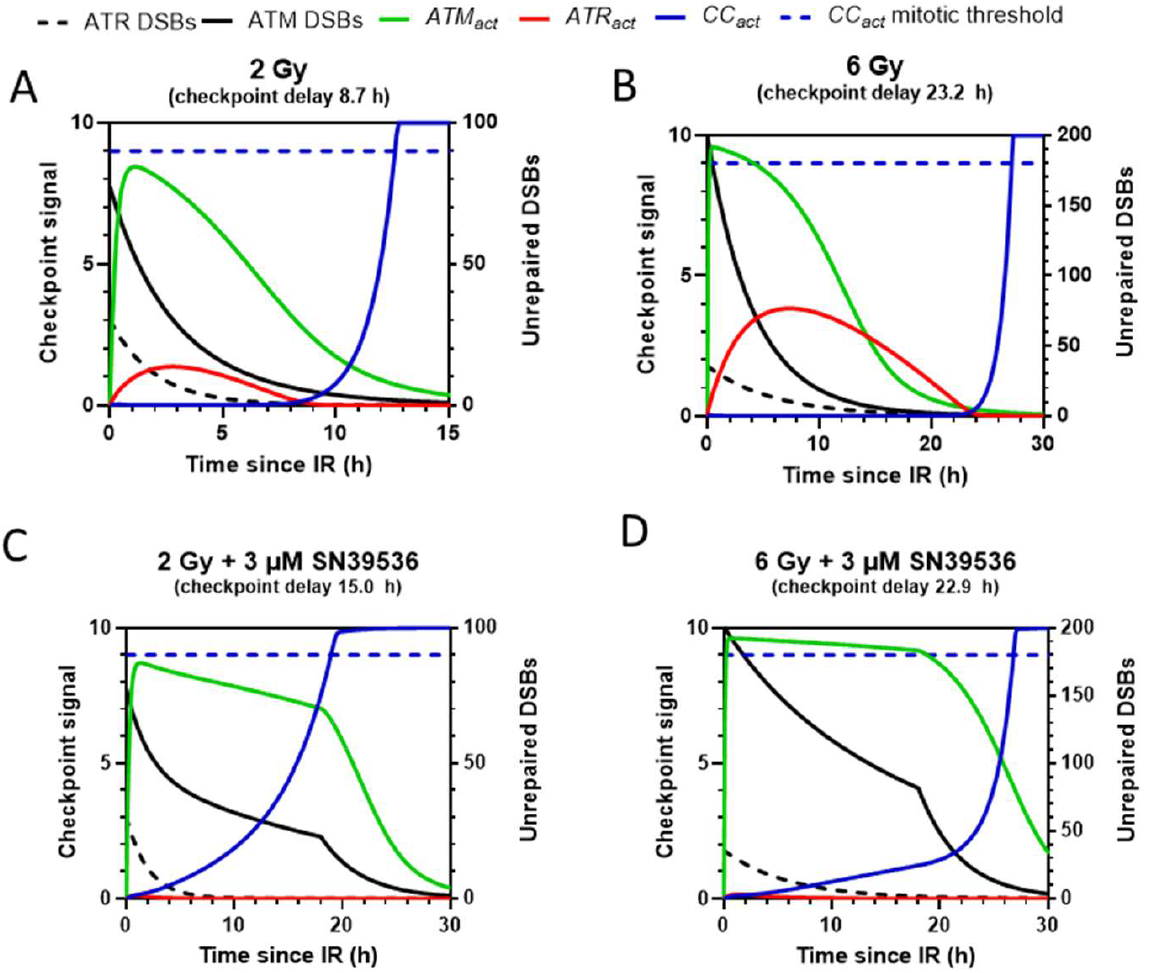
Suppression of cell cycle progression by ATM and ATR signalling, illustrated for cells irradiated at the G2/S border for 2 or 6 Gy IR with or without 3 μM SN39536. In this simulation the DNA-PKi is washed out 18 h after IR. ATM activity (*ATM*_*act*_) is driven by unrepaired DSBs assigned to NHEJ-S and HR (ATM DSBs), and *ATR*_*act*_ from unrepaired DSBs on the HR pathway (ATR DSBs). Both kinases suppress cell cycle progression activity (*CC*_*act*_) through effects on the mitotic kinases (Fig. 2D). The checkpoint delays are the increase in time to mitosis relative to untreated cells for which time to mitosis = 3.9 h.

Checkpoint parameters (Table 2) were estimated by fitting flow cytometry measurements of cell cycle phase distributions after IR with and without 3 μM SN39536 (CCPD dataset; Fig. 8). Experimentally, both 2 and 6 Gy IR alone (Fig. 8A) elicited a marked G2 phase arrest demonstrated by accumulation in G2 phase and an abrupt fall in mitotic fraction as expected for the so-called early G2 checkpoint, which is ATM-dependent and dose-independent ^75^. The block to mitotic entry resulted in a decrease in the G1 fraction and eventually a fall in S-phase fraction. SN39536 alone did not perturb the cell cycle phase distributions while the changes with IR were moderately increased by the DNA-PKi, particularly with 6 Gy with the drug prolonging G2 phase accumulation and M phase suppression (Fig. 8B).

**Fig. 8.**
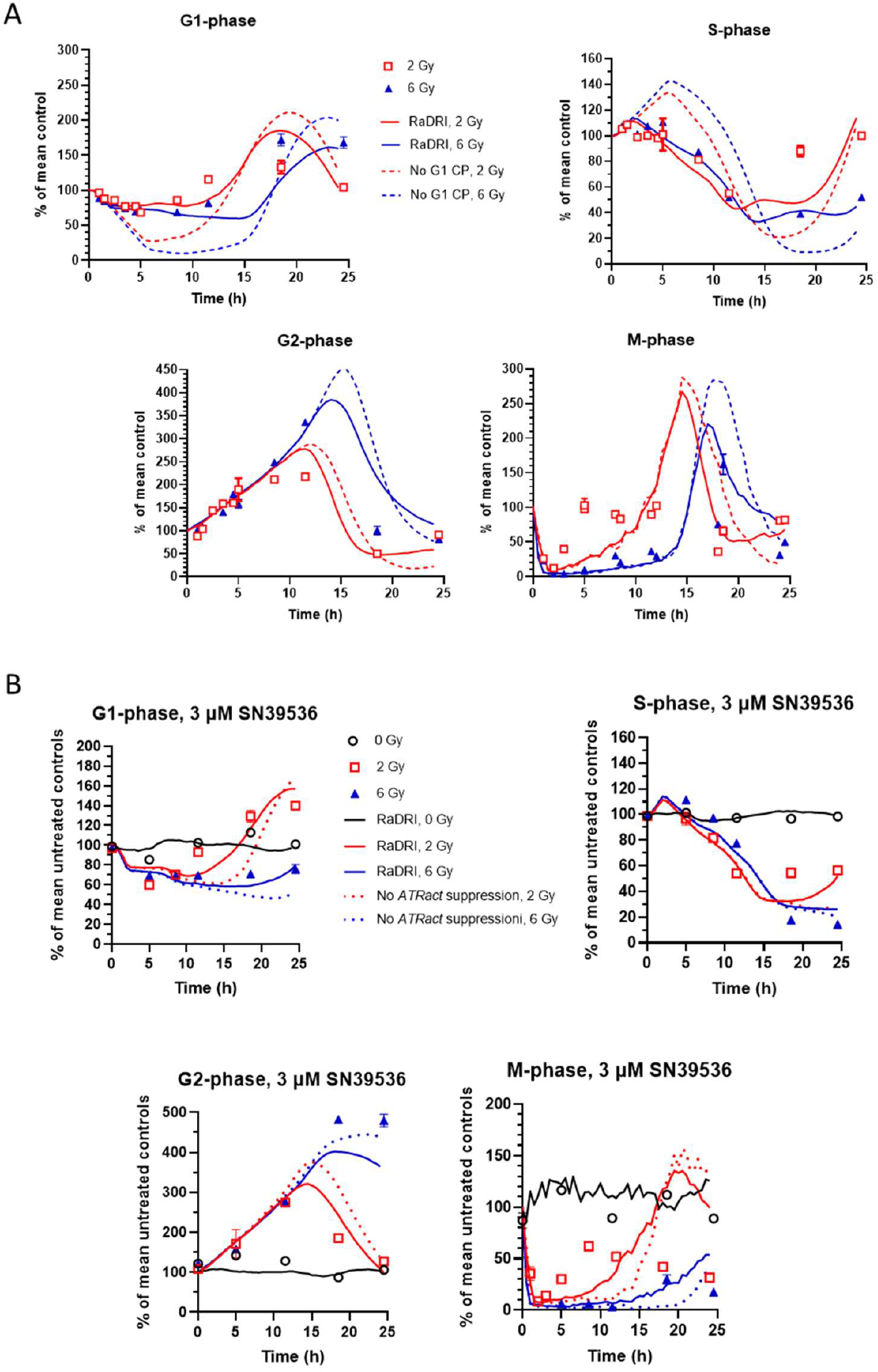
Checkpoint parameter estimation by fitting to flow cytometry data for phase distributions for HCT116/54C cells irradiated (2 or 6 Gy) with or without 3 μM SN39536 (CCPD dataset) at various times after IR. Solid lines are simulations using the parameter values in Tables 1 and 2. Cells were irradiated 24 h after plating 5×10^4^ cells/mL for timepoints < 12 h and 2.5×10^4^ cells/mL for later timepoints, providing < 30% confluency at assay. A: Radiation only. Values (pooled from 3 experiments) are mean and SEM for triplicate cultures in each experiment, expressed as percentage of untreated controls which were averaged across all timepoints. Dashed lines show the G1 fraction predictions in the absence of a G1-phase checkpoint. B: Radiation as for A with 3 μM SN39536, including for 0 Gy, added 1 h before IR and present continuously until assay. Two experiments pooled, with 3 replicates each. Dotted lines are model simulations if *ATRact* signalling is not dependent on DNA-PK catalytic activity.

The combined CCPD dataset was fitted using the G2-phase ATM and ATR-dependent checkpoint model modified from Jaiswal et al.^28^, and an empirical ATM signalling model for the G1- and S-phase checkpoint (Section 3.1.5). Table 2 provides the resulting parameter values, and the sensitivity of the model to parameter variation with respect to the calculated time to next mitosis, when survival is calculated. The model output for CCPD also depends on DSB repair kinetics, controlling ATM and ATR signalling, using parameters determined by fitting CSF (Table 1).

The existence of a G1-phase checkpoint is not immediately apparent in the CCPD data, which does not show accumulation in G1 phase. However, using the model to disable the G1 checkpoint by setting *k*_*atm1G1*_ to zero (dashed lines in Fig. 8A) showed a more severe decline in G1 fraction, with subsequent perturbation in the other phases, confirming the existence of the checkpoint. We also illustrate in Fig 8B the effect of DNA-PK inhibition on the G2 checkpoint; disabling the dependence of *ATRact* on *DNAPKac*t in the model increases the strength of the checkpoint, suppressing the mitotic fraction, because inhibition of DNA-PK by SN39536 no longer compromises ATR activation.

Allowing different G1 checkpoint parameters for daughter cells, which represent the majority of the population in the model at late times, provided a small improvement in the objective function (7.5% decrease in ϕ) with better prediction of phase fractions at 18-24 h with the DNA-PKi. The estimated parameters enforced a surprising increase in cell cycle progression in the daughter cells; we are not aware of a biological mechanism for this, but this small effect made no difference to the predictions of clonogenic surviving fractions.

Originally, we planned to estimate the S-phase checkpoint slowdown, independently of the change in phase distributions, through the reduction in the rate of DNA synthesis (EdU incorporation rate; EDU dataset). Pulse-labelling with EdU for 30 min at various times after 6 Gy (Fig. 9A) demonstrated a modest, transient slowing of DNA synthesis. In combination with 3 µM SN39536 recovery of DNA synthesis rate was delayed, with a small reduction still evident 5 h after IR (Fig. 9B), consistent with extended elevation of *ATMact* from unrepaired NHEJ-S DSBs in the model. However, the reduction in EdU labelling was not well predicted by the S-phase checkpoint parameters estimated in CCPD which showed little recovery by 5 h after 6 Gy (Fig. 9A). The fit could be improved by estimating the ATRact decay parameter *k*_*rd*_ in the EdU experiments which increased its value from 8.8 to 55 and lowered the objective function ϕ for the EDU dataset by 38% (Fig. 9C), but this improvement came at the cost of a 28% increase in ϕ for CCPD. This conflict in fitting the different datasets points to structural limitations in the model. However, below (Fig. 10B) we show that the S-phase slowdown has little impact on the prediction of surviving fractions. In view of this, and the difficulty of accurately measuring the small decreases in EdU incorporation rates, we use the same values for the parameters controlling ATMact across the cell cycle.

**Fig. 9.**
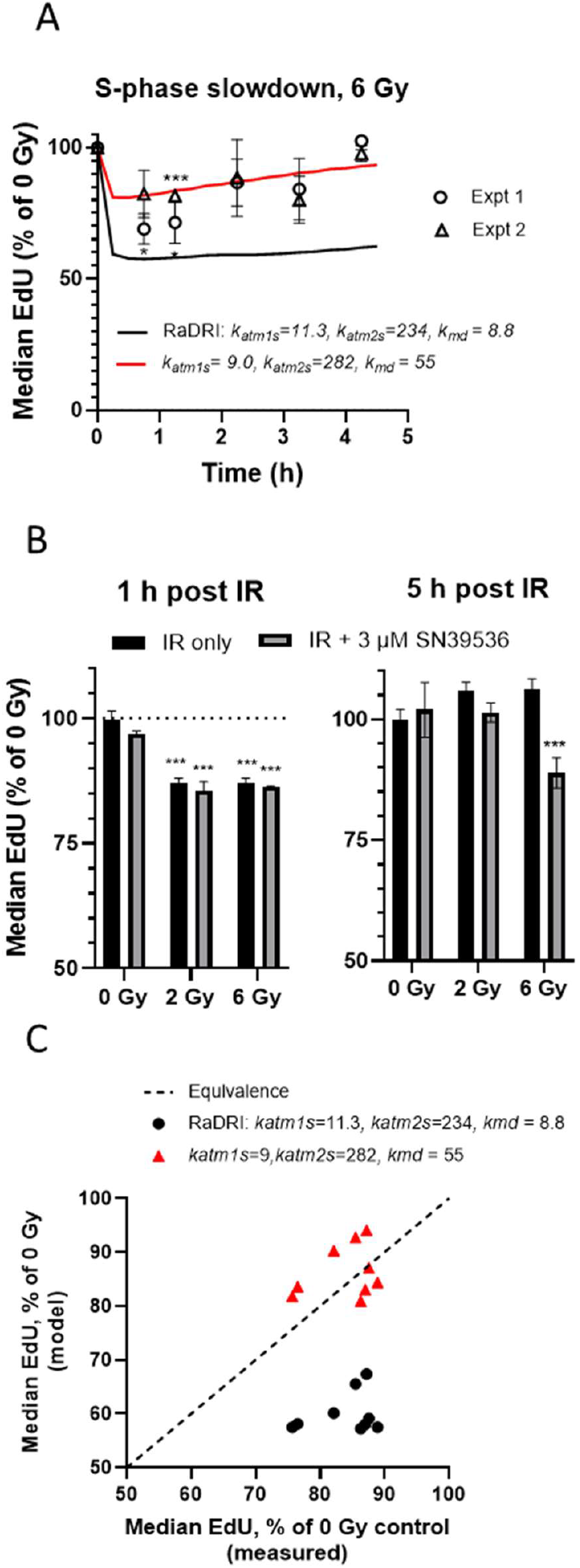
Measurement and modelling of DNA replication rate (EdU incorporation) in HCT116/54C cells (EDU dataset). A: Median EdU fluorescence after 6 Gy irradiation, 20 h after plating 5×10^4^ cells/mL in P60 dishes, as a percentage of untreated controls at each timepoint (at the midpoint of 30 min pulse labelling with EdU). Values are means and SEM for 3 replicate cultures in each experiment. Differences from controls by one-way ANOVA: * p<0.05; **p<0.01. Lines are model fits using the RaDRI parameter set (black) or with three ATM parameters fitted only to the EDU data (red). B: EdU fluorescence as for A, 1 and 5 h after 2 or 6 Gy IR with or without 3 μM SN39536. Mean and SEM are from 3 replicates. ***:p<0.001 relative to zero Gy. C: Model predictions for the EDU dataset (comprising the experimental data in A and B).

**Fig. 10.**
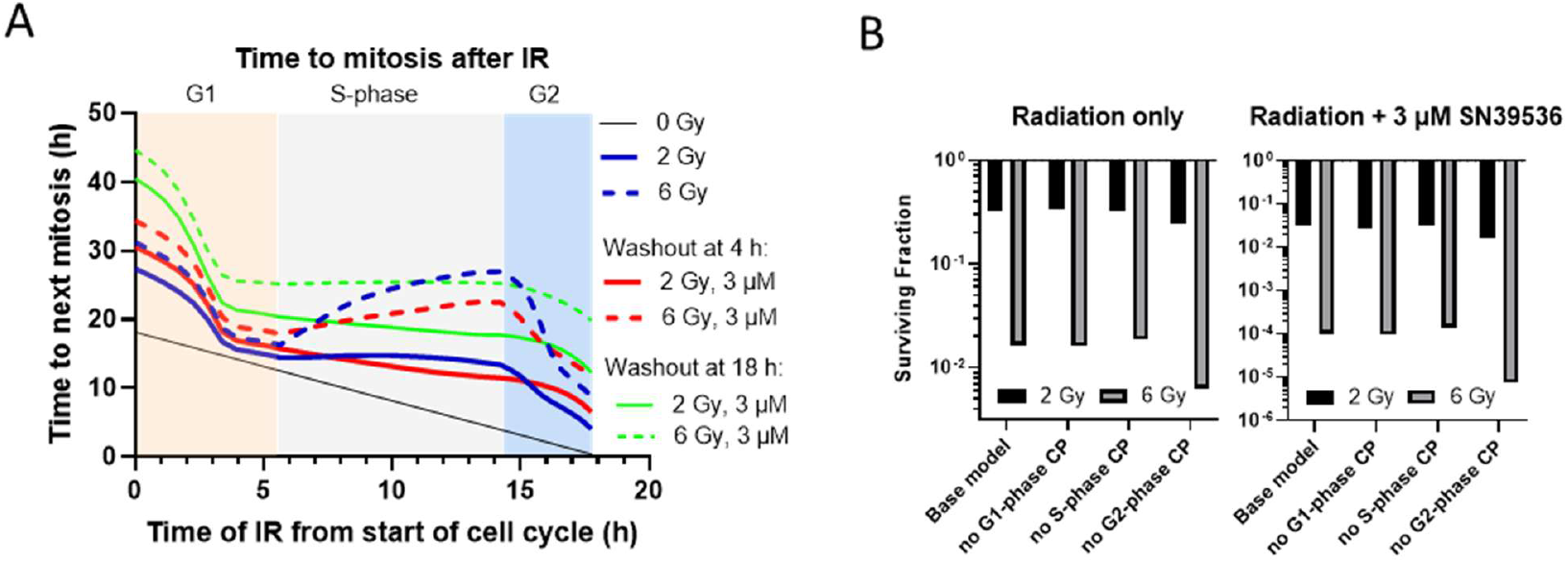
Model simulation of overall checkpoint delay (time to mitosis) and effects on clonogenic cell killing. A: Time to mitosis after 2 or 6 Gy IR alone or following 4 h or 18 h exposure to 3 μM SN39536. 10^4^ cells simulated for IR at each cell cycle phase time. B: Population average clonogenic surviving fraction for 10^4^ log phase cells for the base model, and with each checkpoint individually disabled by setting to zero *k*_*atm1G1*_ for G1 phase, *k*_*atm1S*_ for S phase and for G2 phase a code change to replace the Jaiswal formulation with code for normal phase progression with no slowdown.

The overall effect of the checkpoints on cell cycle delay for cells irradiated at different points in the cell cycle (Fig. 10A) results in large delays for IR in early G1 and late S mid-late S-phase which are exacerbated by SN39536 and increase with duration of drug exposure. (The fall in checkpoint delay for IR later in G1 is a consequence of the delay in the onset of the checkpoint, represented by *T*_*lag*_, and the change in ATM checkpoint parameters between G1 and S-phase). However, by disabling each checkpoint individually in the model, we show using the Table 1 and 2 parameter values that only the G2 checkpoint appreciably influences average clonogenic surviving fraction for the population after IR, either with or without SN39536 (Fig. 10B).

#### 3.4.3 RaDRI modelling of radiation-induced cell killing, with and without SN39536

Clonogenic survival curves for radiation with varying DNA-PKi concentrations and exposure times, and their simulation in RaDRI using the fitted checkpoint and repair parameters (Tables 1 and 2), are shown in Figures 11 and 12 which constitute the CSF (clonogenic surviving fraction) dataset.

**Fig. 11.**
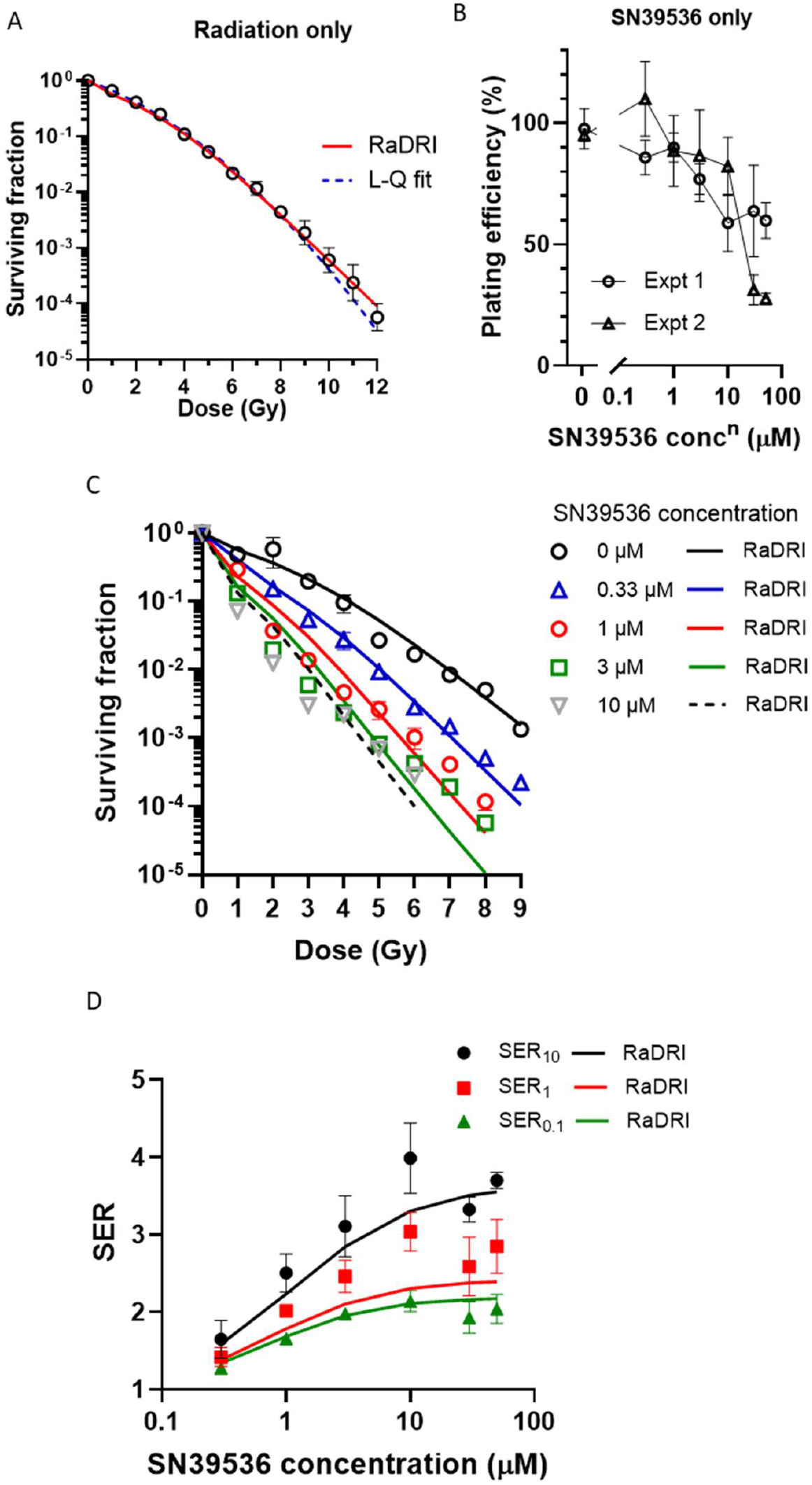
HCT116 **c**lonogenic radiation survival curves ± SN39536, measured 18 h after IR with drug present continuously from 1 h before IR until assay. A: Radiation only. Geometric means and SEM for 5 experiments. Curves show the L-Q and RaDRI fits. B: Cytotoxicity of SN39536 alone. Differences in plating efficiency relative to untreated controls were statistically in Expt 1 at ≥ 10 μM and in Expt 1 at ≥ 3 μM, but over 5 experiments there was no significant difference in plating efficiency between 0 and 3 μM drug (0.84±0.08 vs 0.79±0.06, respectively). C: SN39536 concentration dependence of radiosensitisation. Mean and range for the two experiments in B. D: Drug concentration dependence of SER for 18 h exposure to the DNA-PKi. Points are derived from L-Q fits to the data in C. Lines are the RaDRI model predictions.

**Fig. 12.**
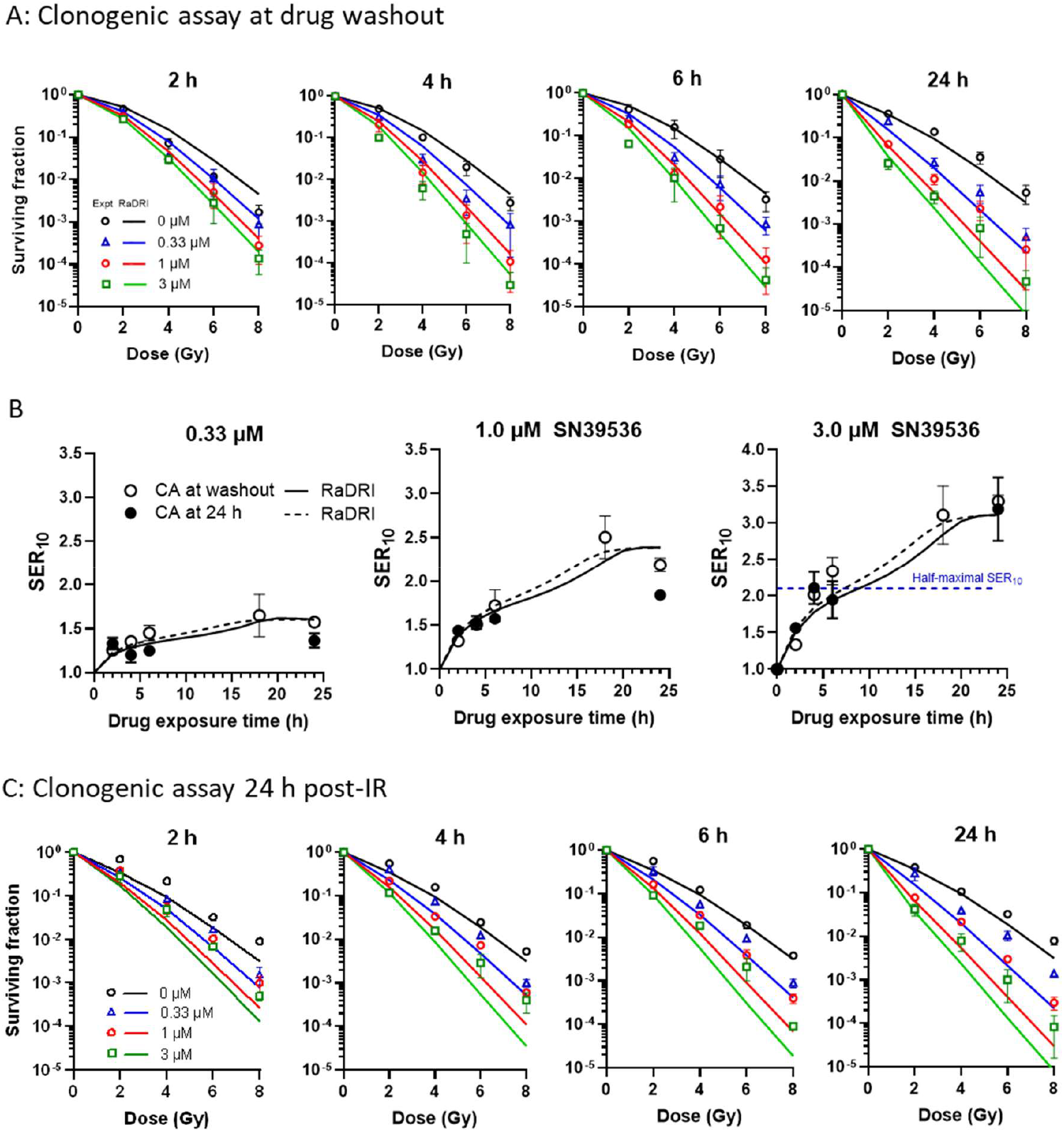
Time and concentration dependence of radiosensitisation of HCT116 cells by SN39536, determined by clonogenic assay. A: Cells were exposed to the drug for 1 h before IR, and until clonogenic assay (CA). Mean and SEM for 3 experiments. B: Dependence of SER_10_ on duration of drug exposure post-IR, and concentration. Symbols are L-Q model estimates (mean and SEM), and line are RaDRI simulations, for washout of the drug at CA or at 24 h. The L-Q estimates for 18 h are re-plotted from Fig. 11C. C: Cells were treated with drug as in B but after removal of the drug after the indicated exposure times the cultures were incubated in fresh medium until CA at 24 h post-IR. Mean and SEM for 2 experiments.

The HCT116 survival curve for radiation alone (Fig. 11A) is fitted equally well by the mechanistic RaDRI model and the empirical linear-quadratic (L-Q) model, with a slight divergence at the highest radiation doses where clonogen survival is both technically and theoretically difficult to define. RaDRI approaches a simple exponential above 6 Gy, rather than the quadratic dependence of the L-Q model, with curvature at lower dose driven mainly by lesion interaction (misjoining) and to a lesser extent by suppression of HR at high dose. SN39535 at concentrations that were minimally cytotoxic alone (Fig. 11B) markedly radiosensitised HCT116 cells exposed continuously to the DNA-PKi until clonogenic assay 18 h after IR (Fig. 11C). Surviving fractions were normalised to the plating efficiency of unirradiated cultures at each drug concentration to correct for any cytotoxicity. The survival curve shape became more linear at high concentration as previously observed with other DNA-PKi or mutant DNA-PKcs ^27,96,97^ which was reflected in the model, although the RaDRI simulations did not precisely capture the survival curve shape at high concentrations and surviving fractions ∼ 1%. As a consequence, SER values were in good agreement with L-Q estimates at 10% and 0.1% survival, but systematically underestimated radiosensitivity at 1% survival (Fig. 11D).

We then evaluated time dependence of radiosensitisation by assaying clonogenic survival 2-24 h after IR (Fig. 12A). Radiosensitisation increased with time, as illustrated by the L-Q model SER_10_ values at all three concentrations tested (Fig 12B); the model provided a good approximation to the survival curves and SER_10_ values. The fits were worst for 2 h drug exposure, with the control survival curve in the model underestimating measured killing, potentially because of perturbation of DSB repair or cell cycle progression when the cells are trypsinised and replated so early after IR. Thus, we also measured radiosensitisation after washing out the drug at various times but not trypsinising for CA until 24 h post-IR (Fig. 12C). The latter experiments were used as a validation set to test the model, rather than for parameter estimation. The two datasets (Fig. 12A,C) provided similar experimental SER_10_ values (Fig. 12B), with the model predictions different slightly because of the daughter cell correction (higher fraction of daughters when CA was delayed until 24 h after IR).

#### 3.4.4 Dissection of mechanisms of radiation-induced cell killing and radiosensitisation by DNA-PKi with RaDRI

Simulation of DNA lesions at mitosis, using the final RaDRI parameters, shows that after IR alone very few radiation-induced DSBs formed in pre-replication DNA survive until mitosis, but that number is increased by 3 µM SN39536 to an extent that depends on when the drug is removed (Fig. 13A). Larger number of DSBs generated in post-replication DNA are present, as a consequence of the shorter time to mitosis, with less dependence on washout time (Fig. 13B). Misjoins at mitosis are most frequent for cells irradiated in G1 (Fig. 13C) because of the time-dependent increase in misjoining in pre-replication DNA, controlled by parameter σ’ in the model (section 3.1.3). The number of misjoins falls in cells irradiated later in G1 because there is less time for σ’ to drive an increase in misjoining before *σ(t)* falls on entry to S-phase. Washout time has little effect because misjoining occurs primarily after DNA-PK inhibition is lifted at washout, and there is ample time for misrepair prior to mitosis whether the drug is removed at 4 or 18 h.

**Fig. 13.**
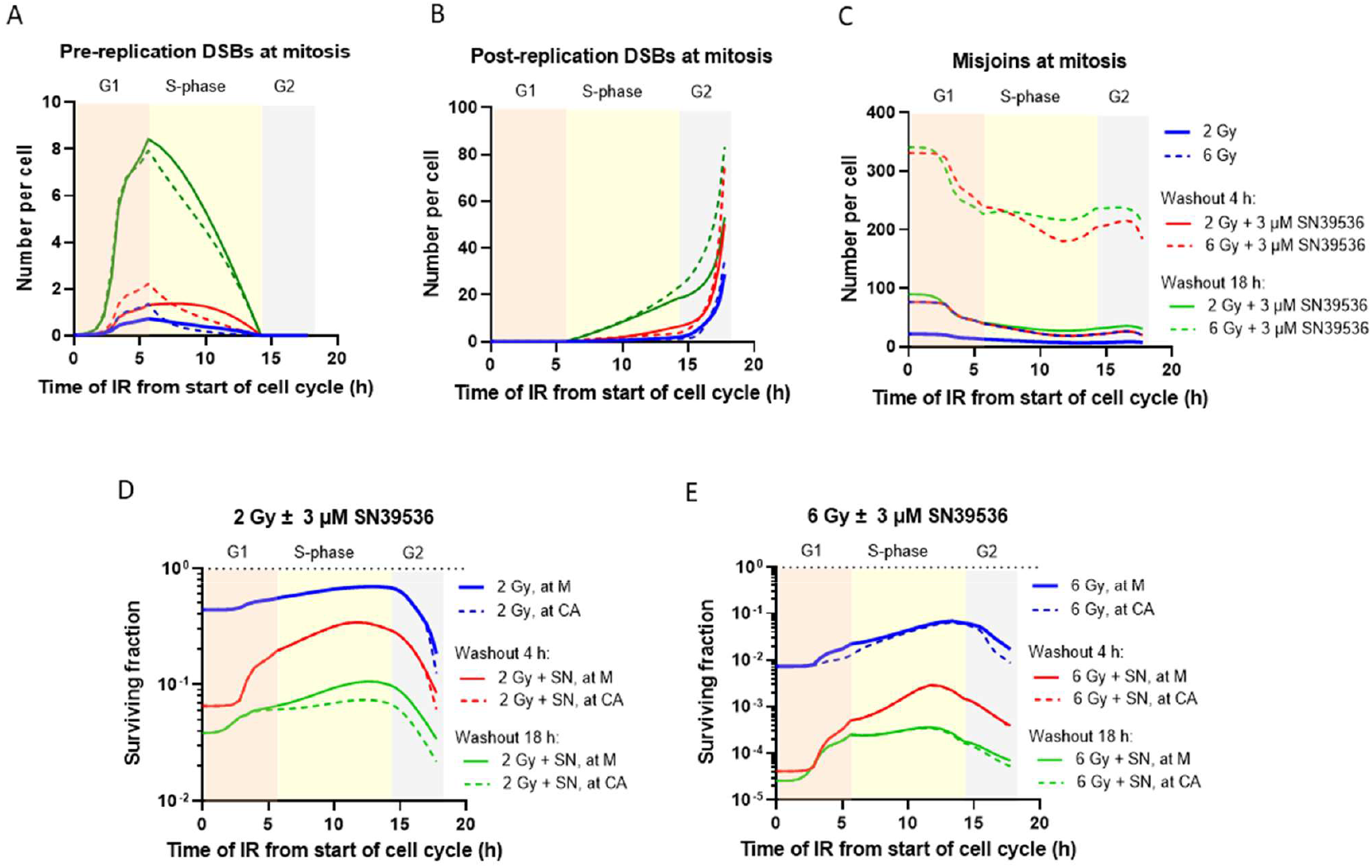
RaDRI simulation of radiation-induced DNA lesions at mitosis, for cells irradiated as different cell cycle stages, with and without 3 μM SN39536, and effect on average surviving fraction of the population. SN39536 is washed out either 4 or 18 h after IR. A-C: Number of DSBs formed in pre-replication DNA, post-replication DNA and misjoins, at mitosis. D,E: Average surviving fraction calculated at mitosis (M, solid lines) and at clonogenic assay (CA; dashed lines) after 2 Gy (D) or 6 Gy (E) IR.

The simulation for cell cycle phase dependence of radiosensitivity at mitosis and clonogenic assay shows the expected increase in radioresistance during S-phase for both 2 Gy (Fig. 13D) and 6 Gy (Fig. 13E) irradiation, which is driven primarily by high-fidelity repair of DSBs in post-replication DNA by HR in the model. However, in the model radioresistance rises prior to entry to S-phase because of the fall in misjoining frequency in late G1 (Fig. 13C). Clonogenic survival is slightly lower for the population at the time of clonogenic assay than the calculated values at mitosis because of the correction for post-mitotic daughter cells (Section 3.1.6). That correction is greatest when survival probability of parent cells at mitosis (*P*_*p*_) is high but also depends on how different the populations are at clonogenic assay relative to at mitosis. Thus, the correction is larger for cells irradiated late in G2 as the population at clonogenic assay then comprises mainly daughters but is less when time to mitosis is highly extended as a result of checkpoint delay (Fig. 10) because the population at clonogenic assay then contains a higher proportion of parent cells.

The contributions of each class of DNA lesion to cell killing was assessed by individually setting the lethality coefficients for pre-replication DSBs (*k*_*dsb1*_), post-replication DSBs (*k*_*dsb2*_), and misjoins (*k*_*mis*_) to zero (Fig. 14). For radiation alone, unrepaired DSBs at mitosis made only a minor contribution to clonogenic cell killing (less for pre-replication DSBs despite *k*_*dsb1*_ > *k*_*dsb2*_; Table 2), while preventing cytotoxicity of chromosomal misjoins dramatically decreased radiosensitivity (Fig. 14A). Initially it seemed surprising that there was no dose response above 1 Gy when killing is only due to DSBs (i.e. when *k*_*mis*_ = 0); we showed that killing by DSBs does increase with dose when the checkpoints are disabled (Fig. 14A), indicating that dose-dependent checkpoint delays counter the increased initial DSB yield at higher dose. The relative contributions of each type of DNA lesion changes substantially with the DNA-PKi; now unrepaired DSBs are the primary drivers of killing at ∼ 1 Gy and post-replication DSBs make a substantial contribution throughout the dose range (Fig. 14B). However, the long checkpoint delays with SN39536 reduce sensitivity to these unrepaired DSBs as shown by the increase in radiosensitivity in the DSB-only model (*k*_*mis*_ = 0) when checkpoints are disabled.

**Fig. 14.**
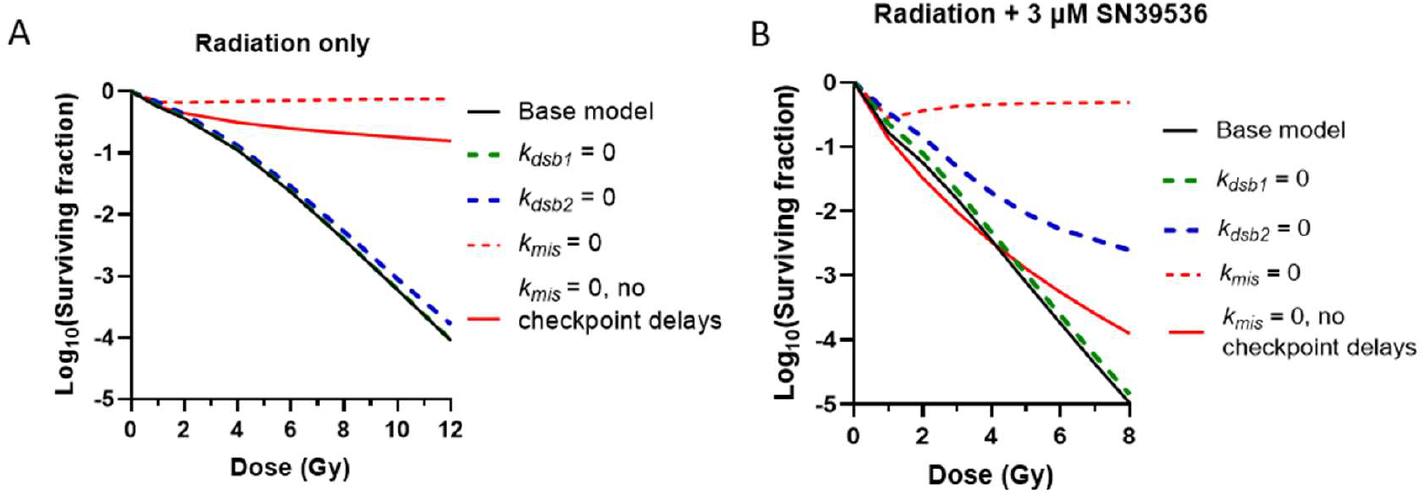
Contributions of DNA lesions to clonogenic cell killing for log-phase HCT116 cells, simulated in RaDRI by setting each of the lethality parameters to zero or with all checkpoints disabled (*k*_*mp1*_ = *k*_*mp2*_ = *k*_*md*_ *= k*_*rp*_ = *k*_*rd*_ = 0) when *k*_*mis*_ = 0. Clonogenic survival was calculated 18 h after IR. A: radiation only. B: radiation + 3 μM SN39536, present continuously until assay.

#### 3.4.5 Pharmacokinetic requirements for radiosensitisation by DNA-PKi

A key objective of this study was to assess time dependence of radiosensitisation by a DNA-PK inhibitor and to represent this using a model formalism suitable for integrating with pharmacokinetic models. Although SN39536 is stable in cell culture, we used RaDRI to simulate radiosensitisation of HCT116 cultures by DNA-PK inhibitors with the same potency at SN39536 but with shorter half-life (Fig. 15). 90% of the maximum additional logs of cell kill with the DNA-PKi is achieved with half-lives of 9.3, 8.7 and 7.7 h at radiation doses of 2, 4 and 6 Gy, respectively (Fig. 15A). 90% of the SER_10_ achieved with a DNA-PK of infinite half-life requires a half-life of 9.2 h, while 50% of the maximal SER is achieved with a half-life of 2.1 h (Fig. 15B). The plasma pharmacokinetic half-life of SN39536 dosed i.p. in mice is ∼ 1.5 h ^97^ suggesting that analogues or dosing routes providing a longer half-life would give greater *in vivo* radiosensitisation.

**Fig. 15.**
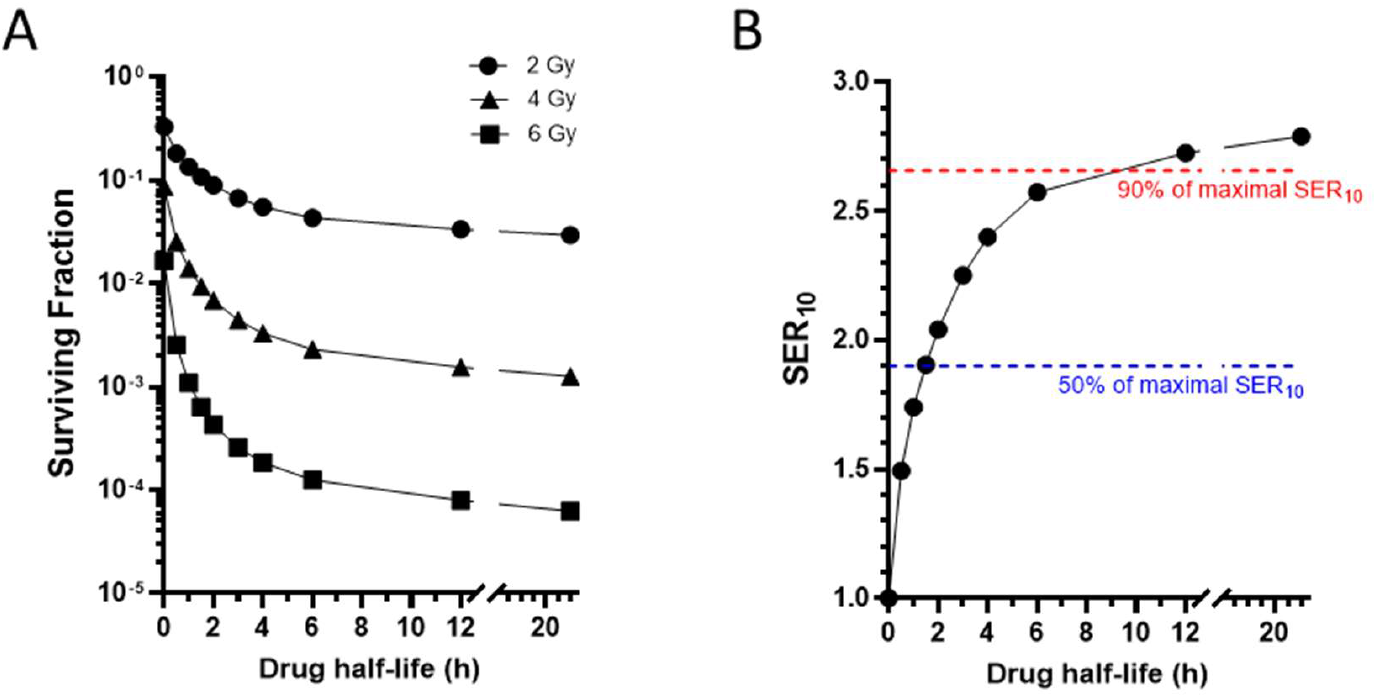
Radiosensitisation of log-phase HCT116 cells *in vitro* by DNA-PK inhibitors at 3 μM, assumed to have the same potency (*EC*_*50*_) as SN39536, as a function of half-life of the compounds. Clonogenic survival is calculated 18 h after irradiation. A: Surviving fraction vs half-life. B: SER_10_ vs half-life.

## 4. Conclusions

The mechanistic RaDRI model provides good prediction of the HCT116 radiation survival curve, and radiosensitisation by a new DNA-PK inhibitor (SN39536), with parameter values estimated by fitting datasets for cell cycle phase distribution and clonogenic cell killing. The model has 7 fitted parameters that determine repair, misrepair and lethality of DNA damage, and 19 fitted parameters describing cell cycle checkpoint delays; together these determine the probability of cell sterilisation which is assessed from the level of DNA damage at mitosis. Additional parameters have been assigned values based on known radiobiology, but this iteration of RaDRI is obviously a highly simplified representation of the complexity of the DNA damage response. Indications of structural inadequacies in the current model include underprediction of SER at low surviving fractions and relatively poor prediction of recovery of the mitotic fraction after IR. Currently, the model neglects features such as potential reassignment of DSBs to alternate repair pathways when NHEJ is inhibited. However, the formalism of the model is readily amenable to extension to additional repair pathways, which will require parameterisation by perturbation with specific inhibitors or genetic modifications.

Despite these limitations, RaDRI offers insights into mechanisms of radiosensitisation by DNA-PK inhibition and interaction with checkpoint delays. The model demonstrates that enhancement of DSB misjoining is a major contributor to radiosensitisation, resulting from the extended lifetime of DSBs when DNA-PK is inhibited coupled with DSB mobility. However, unrepaired DSBs that originate in post-replication DNA also contribute to lethality, much more prominently than with radiation alone. Checkpoint delays have a major protective effect against cell killing by the latter mechanism, suggesting that inhibition of the G2 checkpoint would strongly enhance this component of radiosensitisation as has been demonstrated by synergy between DNA-PKi and ATR inhibition *in vitro* ^98^. We also show that the effect of DNA-PK inhibition on radiation-induced cell cycle perturbation is consistent with the dependence of ATR signalling on DNA-PK catalytic activity; addition of an ATR inhibitor is thus predicted to further enforce premature entry to mitosis when inhibition of DNA-PK is incomplete. Similarly, compromised checkpoint delays may be an important contributor to the reported activity of the DNA-PKi AZD7648 in combination with olaparib against tumour xenografts with mutant ATM ^20,86^.

The expected time dependence of radiosensitisation by DNA-PKi is not immediately obvious as NHEJ is both radioprotective (through reducing unrepaired DSBs at mitosis) and radiosensitising (through enhancement of chromosome aberrations via DSB misjoining). The required drug exposure time is related to NHEJ repair kinetics but not in a straightforward manner. RaDRI accurately models the relationship between DNA-PKi exposure and radiosensitisation, and predicts that half the maximal SER is achieved with approximately 7-8 h exposure (Fig. 15B), while simulating first-order decay of the drug in culture indicates that a pharmacokinetic half-life of approximately 2 h would be required to achieve that level of sensitisation. However, given the interaction with cell cycle progression those relationships will change for tumours which have very different cytokinetics from log-phase cultures *in vitro* ^35^. That question is important because extending exposure to DNA-PK inhibitors beyond the time window defined by DSB repair kinetics risks increasing off-target toxicities.

The present study does not address the key issue of tumour selectivity of DNA-PK inhibitors, or lack thereof ^97,99-101^, which may be influenced in part by those cytokinetic features and needs to be explored further. Our own focus is the development of hypoxia-activated prodrugs to deliver DNA-PK inhibitors selectively to radioresistant (hypoxic) microenvironments in tumours ^102,103^. With that objective, we are currently exploring the use of RaDRI in hybrid continuum (time-dependent Green’s function ^104^)/agent-based (RaDRI) models for mapped tumour microvascular networks to gain insight into the intra-tumour pharmacokinetics and pharmacodynamics of these agents.

## Supporting information

RaDRI clonogenic assay database

## Abbreviations

ATM: ataxia telangiectasis mutated
ATR: ataxia telangiestasis and Rad-3 related protein
CA: clonogenic assay
CMAES_HP: covariance matrix adaptation evolution strategy highly parallelized
DNA-PK: DNA-dependent protein kinase
DSB: double-strand break
EdU: 5-ethynyl-2’-deoxyuridine
HR: homologous recombination repair
IR: ionising radiation
NHEJ: non-homologous end-joining
PEST_HP: parameter estimation highly parallelized
PD: pharmacodynamics
PK: pharmacokinetics
TMEJ: polymerase theta-mediated end-joining.

## Acknowledgments

This work was supported by the Health Research Council of New Zealand (grant number HRC22-444), a Cancer Society of New Zealand Post-Doctoral Fellowship (F.19F) to C.R.H., and a Cancer Society Auckland/Northland Senior Fellowship to M.P.H. The authors would like to thank Dr Lydia Liew for the provision of SN39536 and Ms Way Wong for technical support.

## Supplementary Information

The Fortran code for the RaDRI model is available at Github ^105^. A sensitivity analysis for DNA repair and lethality parameters, and a database of all clonogenic assay experiments in this study, are available as Supplementary Information.

